# Human brain organoids assemble functionally integrated bilateral optic vesicles

**DOI:** 10.1101/2021.03.30.437506

**Authors:** Elke Gabriel, Walid Albanna, Giovanni Pasquini, Anand Ramani, Natasia Josipovic, Aruljothi Mariappan, Friedrich Schinzel, Celeste M. Karch, Guobin Bao, Marco Gottardo, Jürgen Hescheler, Veronica Persico, Silvio O. Rizzoli, Janine Altmüller, Giuliano Callaini, Argyris Papantonis, Olivier Goureau, Volker Busskamp, Toni Schneider, Jay Gopalakrishnan

## Abstract

During embryogenesis, optic vesicles develop from the diencephalon via a complex process of organogenesis. Using iPSC-derived human brain organoids, we attempted to simplify the complexities and demonstrate the formation of forebrain-associated bilateral optic vesicles, cellular diversity, and functionality. Around day thirty, brain organoids could assemble optic vesicles, which progressively develop as visible structures within sixty days. These optic vesicle-containing brain organoids (OVB-Organoids) constitute a developing optic vesicle’s cellular components, including the primitive cornea and lens-like cells, developing photoreceptors, retinal pigment epithelia, axon-like projections, and electrically active neuronal networks. Besides, OVB-Organoids also display synapsin-1, CTIP-positive, myelinated cortical neurons, and microglia. Interestingly, various light intensities could trigger photoreceptor activity of OVB-Organoids, and light sensitivities could be reset after a transient photo bleach blinding. Thus, brain organoids have the intrinsic ability to self-organize forebrain-associated primitive sensory structures in a topographically restricted manner and can allow conducting interorgan interaction studies within a single organoid.

---

3D human brain organoids derived from induced pluripotent stem cells (iPSCs) provide unprecedented opportunities to study the complexity of brain development and diseases (Birey et al., 2017; Gabriel et al., 2016; Lancaster et al., 2013; Mariani et al., 2015) (Anand Ramani, 2020). Brain organoids have helped understanding cellular diversities, complex interactions, and neuronal networks (Birey et al., 2017; Pasca et al., 2015; Quadrato et al., 2017; Xiang et al., 2017). Studies in the 19^th^ century by Pander revealed that during embryogenesis, the retinal anlage develops laterally from the diencephalon of the forebrain, protruding as an optic vesicle (Pander, 1817) (Adelmann, 1966). Huschke then showed that the distal part of the diencephalon invaginates to assemble the optic vesicle (Huschke, 1835) **(Figure 1A)**. In the current century, pioneering works by the Sasai laboratory have demonstrated the remarkable ability of mouse and human pluripotent cells to undergo complex morphogenesis of the retinal anlage that contains cell types derived from neuroectoderm (Eiraku et al., 2011; Nakano et al., 2012). However, *in vivo*, optic vesicles constitute neuroectoderm and surface ectoderm-derived cell types that include specialized neuronal cell types of photosensitive rods and cones, retinal pigment epithelium (RPE), and non-neuronal cell types of lens and cornea (Graw, 2010). Although these structures have been difficult to reconstruct, elegant studies have demonstrated the ability of pluripotent cells to form neuronal cell types of retina individually in the form of retinal organoids (Eldred et al., 2018) (Capowski et al., 2019; Chen et al., 2016; Eiraku and Sasai, 2011; Eiraku et al., 2011; Nakano et al., 2012; Vergara et al., 2017; Zhong et al., 2014) (Cowan et al., 2020).

**Figure 1.**
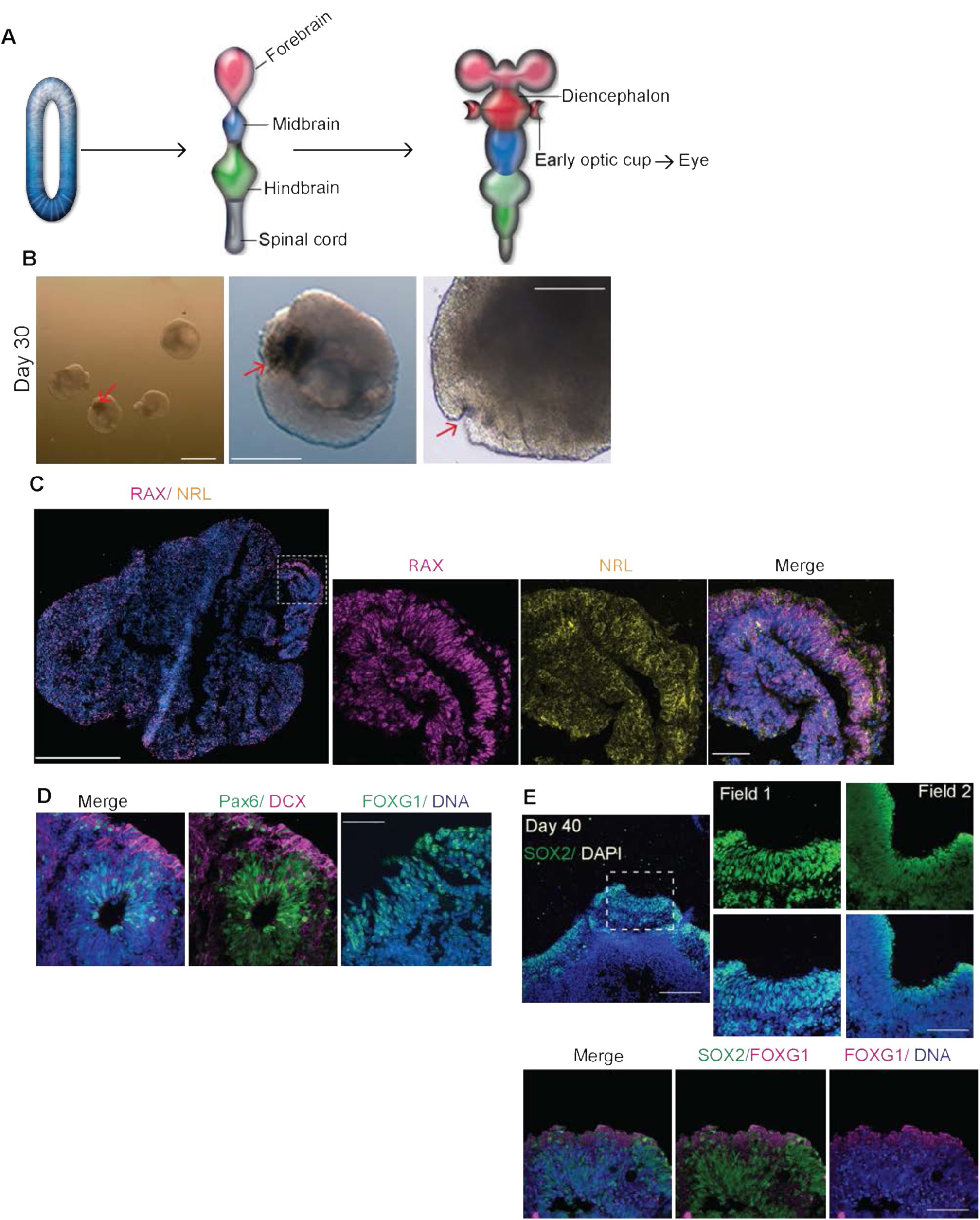
Generation of 3D brain organoids with primordial eye fields from human iPSCs. **A.** Schematic of human embryonic nervous system development. The neural tube (left) segregates into forebrain, midbrain, hindbrain, and spinal cord (middle). Forebrain partially develops into diencephalon from which the early optic vesicles invaginate laterally (right). Optic vesicles later form the eye. **B.** 30-day old organoids are displaying the early occurrence of pigments (red arrows). A red arrow points to an invagination displaying an optic fissure-like structure in the magnified image. Scale bar, 1 mm (left panel) 500 µm (right panel). Representative image, cell line F14536.2. **C.** Organoids display RAX-positive primordial eye field (magenta) and NRL-positive photoreceptor progenitors (yellow). Scale bar, 500 µm for whole organoid and 50 µm (inset). Representative image, cell line IMR-90. Representative images are shown. N=10 organoids from at least 4 independent batches. Representative image, cell line IMR90. **D.** 30-day old organoids show forebrain markers Pax6 (green, left and middle) and FOXG1-(green, right) containing ventricular zones. Scale bar, 50 µm. Representative images are shown. N=21 organoids from at least 4 independent batches. Representative image, cell line IMR-90 and Crx-iPS. **E.** 40-day old organoids show SOX2-positive (green) invaginating regions with FOXG1 (magenta). Scale bar, 200 µm for whole organoid and 50 µm (inset). Representative image, cell line Crx-iPS. Representative images are shown. N=14 organoids from at least 4 independent batches.

Most retinal organoids differentiated from pluripotent cells assemble heterogeneous optic vesicle-like structures, which have to be excised and need further culturing for several weeks. Thus, the isolated retinal organoids differentiated from pluripotent cells do not form *in vivo*-like optic vesicles, cups, or accurately layered RPEs. For example, retinal organoids’ photoreceptors position their outer segments exposed outwardly to the surface, usually capped by RPE cells *in vivo* (Cowan et al., 2020) (Capowski et al., 2019). This could be due to the fact the retinal organoids lack forebrain that is a non-retinal tissue type yet developmentally related to retinogenesis (Capowski et al., 2019).

During embryogenesis, eye development is a complex process orchestrated by a multifactorial repertoire of signaling molecules, receptors, and molecular gradients organized spatiotemporally. Understanding this process will allow underpinning the molecular basis of early retinal diseases. In this regard, it is crucial to study optic vesicles that are the primordium of the eye whose proximal end is attached to the forebrain, essential for proper eye formation. Optic vesicles appear at the 7-8th post-ovulatory week, equivalent to Carnegie stage 18 and 23 of human development (https://embryology.med.unsw.edu.au/embryology/index.php/Carnegie_Stages). A portion of the optic vesicle undergoes an asymmetric self-invagination to produce a “cup” forming a groove. This structure is the optic fissure that closes by the end of the post-ovulatory week 8. Lens vesicle, cornea, and melanin granules (RPE) further continue to appear with retinal differentiation (O’Rahilly, 1975; Oguni et al., 1991). These aspects of embryogenesis have tempted us to challenge whether brain organoids’ self-organization properties can also assemble bilaterally symmetric optic vesicles in a topographically restricted manner mimicking human embryogenesis. If so, the brain organoids could represent biologically relevant 3D tissues.

Brain organoids to assemble functionally integrated optic vesicles allowing inter-organ interactions occurring within a single organoid is intriguing but has not yet been demonstrated. One possibility to generate such hybrid organoids *in vitro* is to fuse distinct cellular origin components of brain and optic vesicles via the recently described ‘assembloid’ approach (Andersen et al., 2020). Interestingly, cerebral organoids could display an immature retina-like structure (Lancaster et al., 2013). Based on these findings, we modified the culturing conditions. We generated human brain organoids with bilaterally symmetric optic vesicles, containing neuronal and non-neuronal cell types and exhibiting functional circuitry. Importantly, we could generate these organoids within 60 days, a time frame that parallels with human embryonic retina development and therefore feasible to conduct multiple *in vitro* experiments (O’Rahilly, 1975).

## Results

### Generation of 3D brain organoids with primordial eye fields

We previously described a protocol of inducing differentiation into neural epithelium directly from iPSCs (Gabriel and Gopalakrishnan, 2017; Gabriel et al., 2017; Gabriel et al., 2016) (Anand Ramani, 2020). These brain organoids expressed retina and eye-related genes (data not shown) but never developed into visible optic vesicles. We, therefore, modified our protocol starting with a low-density cell number (1x 10^4^ iPSCs as starting number for one organoid). Our intention was not to force the development of purely neural cell types at the earliest stages of organoid differentiation. Therefore, we provided retinol acetate ranging from 0 to 120 nM to the culture medium at a time point when the neuroectoderm expands (for detailed information see the Materials and Methods section). Retinoic acid is critical for early eye development as it is a paracrine inhibitor of the mesenchyme around the optic cup (Rosen and Mahabadi, 2020) (Cvekl and Wang, 2009) (Janesick et al., 2015). An addition of 60 nM of retinol acetate reproducibly induced the formation of pigmented structures, possibly primordial eye fields at around day 30 **(Figure 1B).** Interestingly, a region proximal to the pigmented area often displayed an invagination suggesting an optic vesicle’s assembly in a brain organoid. By contrast, organoids differentiated from iPSCs reprogrammed from adult retinal müller glial cells generated strongly pigmented structures within 20 days (Slembrouck-Brec et al., 2019) **(Figure S1A).** Pigmented areas restricted to one pole of the organoid suggested the presence of the forebrain-like region where the primordial eye field develops.

Next, we tested for the presence of eye field patterning markers in these organoids. Immunostaining for identity markers revealed RAX, NRL, Pax6, and FOXG1-positive progenitor cells in this region **(Figure 1C and D).** The primordial eye field, which arises in an area lateral to the diencephalon, is enriched with RAX, a transcription factor essential for eye field patterning and to define optic vesicles developing from the diencephalon (Furukawa et al., 1997a) (Mathers et al., 1997; Stigloher et al., 2006). Likewise, the transcription factor NRL specifies the cell fate of rod photoreceptors (Kim et al., 2016). FOXG1 is initially expressed in the prosencephalic neuroepithelium and later involves in the telencephalon and eye field segregation (Stigloher et al., 2006). Around day-40 organoids, we observed prominent SOX2-positive invaginating regions that displayed gradients of FOXG1, suggesting that the optic field in these organoids is segregated to the forebrain region. Thus, in this method, it appears that the retinogenesis is restricted at one side of the organoid as specified by the progenitor markers VSX/SOX2 **(Figure 1E and S1B).**

To investigate the cell diversity of primitive eye field containing brain organoids, we performed single cell RNA-sequencing (scRNA-seq) of 30-day old organoids. Out of two sequencing batches of pooled organoids, the transcriptomes of 3511 single cells have been analyzed. Embedding the cells in a uniform manifold approximation and projection (UMAP) revealed the presence of a group of cycling progenitor cells developing towards multiple neuronal cell fates. Cluster analysis and marker genes detection revealed eight main transcriptionally distinct cell populations (Clusters C1 to C8) **(Figure 2A-C and Table 1).** The organoids contained cell clusters of neural progenitors, forebrain development and neurons (C1 to C4). We also identified cell clusters of developing optic vesicles (C6) and pre-optic area (C7) which were segregated from neuronal cell types **(Figure 2A-C).** Four telencephalic clusters containing radial glial (as defined by SOX2 expression), cycling cells (as defined by AURKB), developing forebrain (as defined by LHX2 and LHX9) and neuronal (as defined by DCX, NCAM1 and MAP2) cell classes which occupied a major proportion of total cells (Rétaux et al., 1999) (Peukert et al., 2011) (**Figure 2A-B).** Our analysis also identified a glial cell cluster (C3) of which a fraction of cells expresses markers specific for müller glia (S100A16, APOE, ITM2B, and COL9A1) and microglia (ICAM1 and AIF1). Both cell types are required at the early stages of retina development, such as regulation of retinal ganglion cell subsets and synapse pruning (Li et al., 2019) **(Figure 2B).**

**Figure 2.**
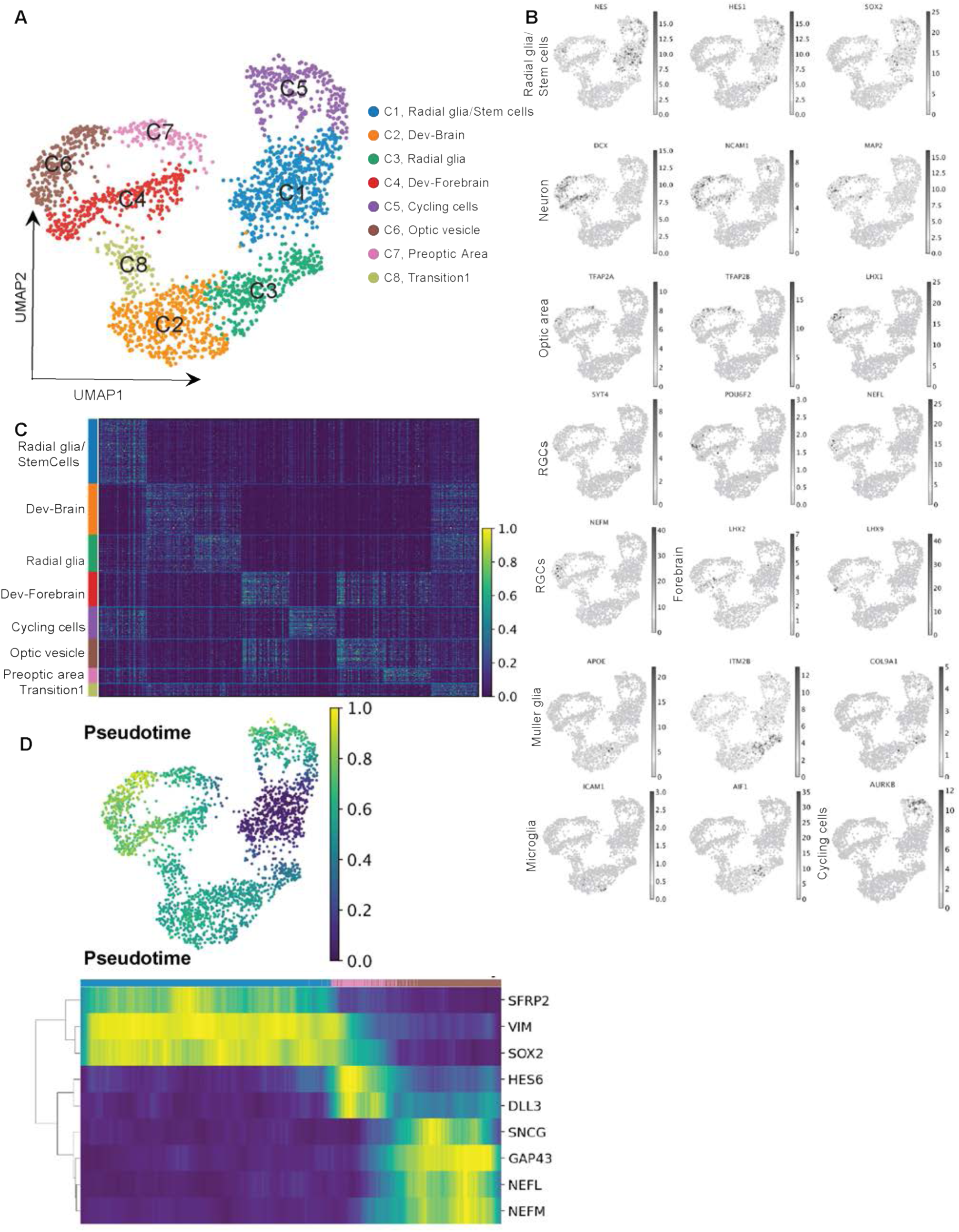
scRNA sequencing of 30-day old brain organoids containing primitive eye fields. **A.** UMAP plot showing cells from 30-day old organoids clustered in eight groups based on transcriptome similarity. Major cell types clustered into eight distinct groups. Legends at the right name the clusters. Besides a pool of glia/stem cell-like cells, we observe cycling cells and trajectories of developing brain and early retina-related cell types. **B.** UMAPs showing marker genes used for annotation of the clusters. **C.** Heatmap showing the expression of the top 20 markers calculated by Student-t-test for each cluster. See Table 1 for the complete list of markers. **D.** Upper Panel: Pseudotime trajectory showing radial glia/stem cells as root (blue cells), and all the other cells colored according to the computed pseudotime (green to yellow). Lower panel: The heatmap shows the expression of key genes identified by Sridhar et.al. 2020 for retinal ganglion cells formation in the early developmental steps of the retina (Sridhar et al., 2020). Heatmap shows the expression of such genes in the cells belonging to C1, C7, and C6, sorting them according to the pseudotime. Plotted expression values are convolved in each row by a window of 20.

**Table 1:** Related to main figure 2. Differentially expressed genes used to define cell clusters of early brain organoids with primitive eye fields.

Further investigation of marker genes revealed the presence of two more clusters of developing optic vesicle (C6) and a pre-optic region (C7) expressing early transcription factors TFAP2A/B and LIM homeobox gene LHX1 essential for optic fissure closure and precise timing of neural retina differentiation (**Figure 2A-B).** TFAP2A expression has also been detected in the lens and neural retina and mutations in TFAP2A is associated with branchio-oculo-facial syndrome (BOFS) in which its associated structures are defective (Min et al., 2020). On the other hand, LHX1 is expressed in the forebrain at a time point that parallels to the formation of neural retina (Inoue et al., 2013). The optic vesicle cluster (C6) also contained cells that strongly expressed NEFM and NEFL suggestive of developing retinal ganglion cells **(Figure 2B)**. Furthermore, by computing pseudotime in our dataset, we performed a trajectory analysis of the cells developing towards the pre-optic cluster. Interestingly, in 30-day old brain organoids containing primitive eye fields express essential marker genes of developing retinal ganglion cells with the same pattern of appearance, as previously described in the human fetal retina (Sridhar et al., 2020) **(Figure 2D)**. Overall, our combined data demonstrate that *in vitro*-derived brain organoids with developing optic vesicle recapitulates both brain and optic-related cell composition and transcriptomic signatures.

### Brain organoids progressively develop optic vesicles over time and reveal brain, early retina and eye-related cell populations

Continued culturing of brain organoids resulted in the progressive development of these pigmented regions forming one or two intensely pigmented optic vesicle-like structures between days 50 and 60. We term these organoids as optic vesicle brain organoids “OVB-Organoids” **(Figure 3A-C).** This method is robust and reproducible because we could generate OBV-Organoids across five independent iPSC donors **(Figure 3B).** For example, 86 out of 95 organoids from the IMR-90 iPSC line could develop easily recognizable bilateral symmetric optic vesicles with an overall success rate of 226 (66%) (**see Table 2 for details**). None of the organoids derived from any of the five tested iPSC donors formed more than two pigmented regions. Intriguingly, these pigmented optic vesicles were exclusively restricted to one pole of the organoids near each other, suggesting an area topographically patterned at the forebrain region.

**Figure 3.**
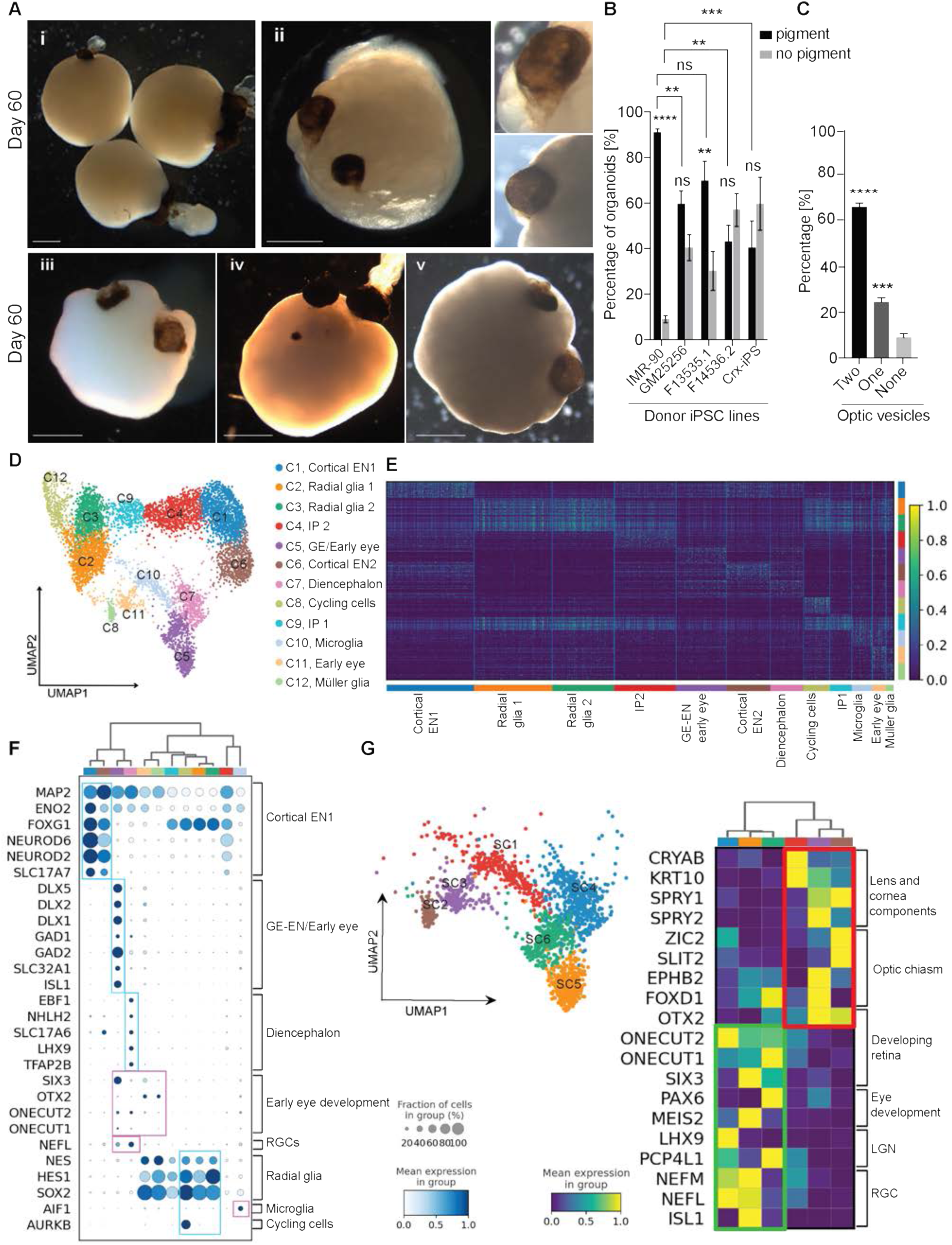
OVB-Organoids reveal brain, early retina and eye-related cell populations. **A.** Macroscopic views of 60-day-old organoids with each containing bilaterally symmetric pigmented optic vesicles **(i-v).** Insets of **(ii)** show close up view of individual pigmented regions. Scale bar, 1 mm. Representative images from at least four independent batches of organoids. N≥32 organoids from ≥3 independent batches. Cell lines IMR-90 **(i)** + **(ii),** F13535.1 **(iii + v),** GM25256 **(iv).** **B.** Bar diagram quantifies organoids with pigmented optic vesicles. On an average, iPSC line of donor 1 (IMR90) yields the highest number organoids with pigmented optic cups (91%). Donor 2, 3, 4 and 5 yield 65%, 70%, 43%, and 40% respectively. For each cell line, at least three different batches were generated. Donor 1: number of organoids = 95, n = 5 batches; donor 2: number of organoids = 280, n = 5 batches; donor 3: number of organoids = 87, n = 4 batches; donor 4: number of organoids = 32, n = 3 batches. donor 5: number of organoids = 241, n = 3 batches. Donor1 to 5 are IMR-90, GM25256, F13535.1, F14536.2 and Crx-iPS. Statistical analysis within and across cell lines. Two-way ANOVA followed by Sidak’s multiple comparisons test. Significance within cell lines: IMR-90: n = 5, p****<0.0001; GM25256: n = 5, p = 0.7359; F13535.1: n = 4, p = 0.1852; F14536.2: n = 3, p > 0.9999; Crx-iPS, hiPSC: n = 3, p > 0.9999. **C.** Bar diagram quantifies percentages of two-, one- or no optic vesicles-containing organoids derived from donor 1 (IMR90) iPSC cells. In an average approx. 66% of organoids differentiated from this cell line yielded two pigmented optic vesicles restricted at one pole. N = 95 organoids from 5 batches. **D.** UMAP plot showing major cell types of OBV-Organoids clustered into twelve distinct groups according to their transcriptome. Both brain and retina-related identity of cells have been found: Radial Glia, intermediate progenitors (IP), cortical excitatory neurons (EN), diencephalon neurons, ganglion eminence (GE), early eye-expressing markers cells, microglia, and Müller Glia. **E.** Heatmap highlighting the expression of the top 20 marker genes for each cluster computed by Student-t-test. See Table 3 for the complete list of marker genes. **F.** Dot plot showing mean expression in each group and fraction of cells expressing sets of literature-based marker genes. Cortical EN, GE-EN and Diencephalon genes were obtained by Kanton et.al. *2019* (Kanton et al., 2019). The plot shows brain-related clusters expressing higher levels of key brain marker genes (light blue boxes). Simultaneously, cluster C5, C7, C8, C10 and C11 show an additional expression of retina development markers or microglia (pink boxes). **G.** Subclustering clusters expressing some retinal-related marker in the dataset (see pink boxes at 3F). The UMAP shows the six subclusters. In the matrix plot, we plotted a group of genes related to specific areas of the eye or cell type: lens and cornea components, optic chiasm, developing retina, eye development, lateral geniculate nucleus (LGN), and retinal ganglion cells (RGC) (Kanton et al., 2019).

**Table 2:** Statistical summary of organoids and their usage in different figures in this manuscript.

**Table 3:** Related to main figure 2. Differentially expressed genes used to define cell clusters of OVB-Organoids

To analyze the cell diversity of OVB-Organoids, we analyzed a total of 7298 high-quality single cells out of six organoids differentiated from IMR-90. Clustering analysis resulted in twelve cluster identities (Clusters C1 to C12) showing distinct transcriptome profiles **(Figure 3D, E and Table 3).** UMAP embedding showed the presence of a main group of cells expressing pan-neuronal markers of DCX and MAP2, telencephalon marker EMX1 (upper part of the UMAP), and a smaller subset of cells expressing diencephalon marker DLX2 (C5, C7). To further annotate the neuronal identity of these cells, we used a set of previously described markers expressed in different parts of brain organoids (Kanton et al., 2019). Cortical excitatory neurons (EN) were identified by the expression of canonical marker MAP2, neurogenic factors of NEUROD 2 and 6, glutamatergic neurons (SLC17A7), ENO2 containing mature neuronal types including FOXG1, a critical transcription factor for forebrain development (Hettige and Ernst, 2019) **(Figure 3F)**. The ganglionic eminence (GE) appears during the embryonic nervous system development, ensuring the proper guidance of cell and axon migration (Nery et al., 2002) (Métin and Godement, 1996). The cluster GE-EN early eye development (C5) comprises cells with GE’s signatures and GABAergic signatures (for example, identity markers of DLX1, 2, 5, and ISL1), suggesting that the OVB-organoids are still at the early phase of development. The significant proportion of cells in this cluster expressed endogenous GAD2 that is expressed in the intrinsic photosensitive retinal ganglionic cells (ipRGCs) (Sonoda et al., 2020). The diencephalon cluster (C7) comprises cells with signatures of diencephalon developmental genes such as LHX9, EBF1, and NHLH2.

Perhaps the most intriguing clusters are early eye development (C5 and C11), diencephalon (C7), müller glia (C11), and RGC clusters that exhibit clear eye development signatures enriched with cells expressing SIX3; a transcription factor plays a crucial role in eye formation in the forebrain (Loosli et al., 1999) (Carl et al., 2002). These clusters also contain OTX2-expressing cells, which activates the downstream transcription factors of PRDM11 and VSX2 to determine photoreceptor and bipolar cell fates (Goodson et al., 2020) (Ghinia Tegla et al., 2020). These cell clusters also harbor factors that regulate the development of horizontal cells, ganglion cells, cones, and amacrine cells (for example, ONECUT1/2 and NEFL). Cells in these clusters also abundantly express progenitor markers of FOXG1, SOX2, and HES1. Among these, HES1 is a transcriptional repressor critical for the timing of retinal neurogenesis **(Figure 3F)** (Lee et al., 2005). HES1 is required for the differentiation and morphogenesis of early lens, optic vesicle, and RPE, as this molecule is localized broadly in these tissue types. The C5 and C11 clusters also contained signatures of preplacodal ectoderm (PPE) and lens ectoderm cell clusters (DLX5, 6, BMP7, MAB21L1, MEIS1 and 2) **(Figure S2).** The PPE contains both neurogenic and non-neurogenic placodes, which give rise to ocular surface ectoderm (Lleras-Forero and Streit, 2012), suggesting that the OVB-Organoids could include non-neuronal cell types of optic vesicles such as the lens and cornea whose biogenesis begins from the surface ectoderm.

Thus, to further delineate the degree of cellular diversity generated within these OBV-organoids, we more closely dissected clusters exhibiting retinal-related expression markers in **Figure 3F** (C5, C7, C8, C10 and C11) by a second iteration of clustering (SC1 to SC6) **(Figure 3G and S3)**. We then wanted to identify cells expressing signatures of lens and cornea components (SC1), optic chiasm (SC2 and SC3), developing retina, eye, lateral geniculate nucleus (LGN), and RGCs (SC4 to SC6). Examining these subclusters, we noticed CRYAB and KRT10 expressing cells in a distinct subcluster (Color code: Red, SC1). These genes signify the formation of the mammalian lens and corneal epithelium. Similarly, we noticed SPRY1 and 2 (lens and corneal marker), essential for eyelid closure (Purple and brown, SC2 and SC3). Cells within this subcluster express ZIC2 and SLIT2 (Brown, SC2), transcription factors determine the routing of RGC axons at the optic chiasm to respective hemispheres, a patterning event that is critical for binocular vision (Lee et al., 2008) (Herrera et al., 2003). Other closely related subclusters (SC4 to SC6) contained cells expressing signatures of developing retina (Turquoise; ONECUT1, 2 and SIX3), eye development (Orange and green; PAX6 and MEIS2), and RGCs (Turquoise and orange; NEFM, NEFL, and ISL1). Intriguingly, within this subcluster, we also identified cells expressing LHX9 and PCP4, molecules critical for forming lateral geniculate nuclei, primary sensory thalamic nuclei receive a major sensory input from the retina (Iwai and Kawasaki, 2009). Hierarchical clustering based on the subset of genes shows a main distinction between SC1-3 and SC4-6. Together, these analyses indicate that distinct neuronal and non-neuronal cell types in OBV-Organoids transcriptionally resemble the endogenous counterparts.

### Comparative transcriptome of OVB-Organoids reveals diverse retinal cell types

Given the relatively small numbers of cells in each subcluster sampled from OVB-Organoids, it is possible that our scRNA analysis overestimated the cell diversity. To enhance the coverage and to further delineate differential gene expression, and cellular diversity, we generated and compared transcriptome profiles of early brain organoids with primordial eye fields to those of OVB-organoids using bulk mRNA sequencing **(Table 4).** Following data processing, read mapping and variance normalization, principal component analysis (PCA) showed two well-defined subgroups contributing up to ∼80% of observed sample distance corresponding to OVB-Organoids and early brain organoids. Differential gene expression analysis revealed ∼4,000 genes with significantly changing mRNA levels in OBV-Organoids compared to early brain organoids (q-value <0.05, **Figure S4**). Of these, ∼3/4 were up regulated and most probably signifying the activation of the developmental pathway leading to optic vesicle formation. This interpretation was substantiated by affinity propagation clustering to identify gene signatures from differentially expressed genes resulting in seventy-five sub-clusters further grouped into nine super-clusters displaying distinct molecular pathways. In addition to synapse maturation, myelination, cell proliferation, and cell cycle regulation, which all signify neural tissue morphogenesis and differentiation, we also observed up-regulation of genes involved in the detection of light stimuli, in visual perception, as well as in compound eye morphogenesis (super-clusters 4 and 7, respectively; **Figure S5A**).

**Table 4:** Bulk mRNA sequencing data comparing transcriptome profiles of early brain organoids with primordial eye fields to those of OVB-organoids.

Gene set enrichment analysis (GSEA) of super-cluster 4 that carries the visual perception-related signature coupled to leading-edge analysis, revealed an array of genes associated with early retina development, including early retinal progenitor, retinal ganglion markers, lens as well as transcription factors of photoreceptor cell (e.g., *RPE65*, *SFRP5*, *FGF19*, *CRX*, *RCVRN*, *RAX*, *VSX2*, *LIN28B*, *PRTG*, *ATOH7*, *DLX2* or *POU4F2*; **Figure S5B**). Furthermore, direct comparison to transcriptomic data from fetal retina showed that there indeed exists an apparent correlation between the fetal retina and OVB-Organoids in terms of expressed genes relevant to horizontal, amacrine, bipolar, Müller glia, progenitor and retinal ganglion cells (**Figure S6**).

Trying to find out where OVB-Organoids fit in the time scale of embryonic development, we compared our RNA-seq data to the collection of gene expression data provided by the LifeMap Embryonic Development & Stem Cell Compendium(Edgar et al., 2013). This comparison revealed that OVB-Organoids are enriched for genes relevant to compound eye development, retinal recycling, phototransduction, including cell types derived from surface ectoderm giving rise to the lens and cornea (e.g., *CRYAB*, *CRYBB3*, and *OPTC*; **Figure S7**). In summary, most aspects of our transcriptomic data are in strong correlation with the development of a neural retina in the optic vesicles of OVB-Organoids.

### OVB-Organoids display immature neural retina

To validate our transcriptomic data, we characterized the neural retina by immunostaining for known markers in organoid sections transversely cut through an optic vesicle. The pigmented RPE layer has strongly interfered and was problematic with the immunostaining and the visualization of nuclei. Therefore, we either used sections briefly treated with 0.5 % hydrogen peroxide, which reduces the pigment intensity, or sections where RPE positioning did not hinder the visualization. In these sections, we detected OTX2-positive nuclei adjacent to RPE cells forming a layered structure. OTX2 that critically determines photoreceptor cell fate consistently present at one side of the organoid restricted to RPE layer helped us coordinating our imaging experiments. On a few occasions, OTX2-positive nuclei were also positive for ONECUT2 **(Figure 4A-B and S8A).** These data indicate progenitors’ presence that can promote the fates of both cone photoreceptors and horizontal cells(Emerson et al., 2013).

**Figure 4.**
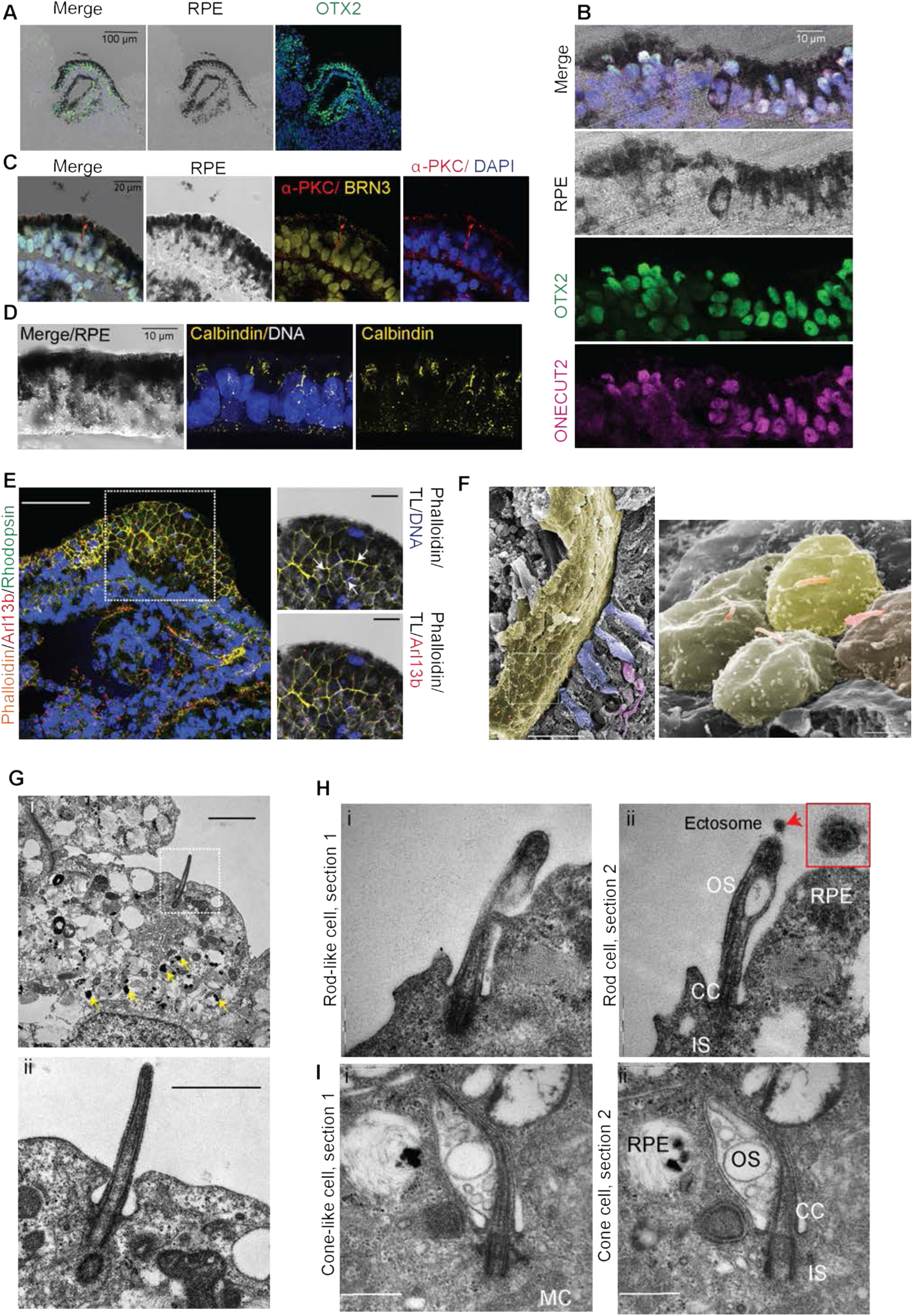
OVB-Organoids display developing neural retina. **A.** Transverse section through an optic vesicle of an organoid shows RPE cell layer imaged by transmission light (Dark pigmented). OTX2-positive nuclei (green) are seen adjacent to RPE layer cells forming a layered structure. Scale bar, 100 µm. N=15 organoids from at least 3 independent batches. Representative image, cell line GM25256 and IMR90. **B.** OTX2-positive (green) nuclei are also positive for ONECUT2 (magenta). RPE layer is imaged by bright fields. Scale bar, 10 µm. N=15 organoids from at least 3 independent batches. Representative image, cell line GM25256 and IMR90. **C-D.** Transverse section through an optic vesicle. Beneath RPE cell layer BRN3-positive RGCs are present. α-PKC-positive (red) bipolar cells and calbindin-positive cells (yellow) (**D**) are infrequently present in these layers. Note the presence of intensely pigmented RPEs and dark melanin pigments that interfere with staining and imaging **(D).** Scale bar, 20 µm for **C** and 10 µm. N=15 organoids from at least 3 independent batches. Representative image, cell line GM25256 and IMR90. **E.** Section through the RPE cell layer of an organoid. The dotted box marks the pigmented area consisting of honeycomb shaped RPE cells visualized by Phalloidin (yellow, arrows) and primary cilia (Arl13B, red). Retinal pigment (black) hinders the visualization of nuclei (blue) and Rhodopsin (green). Scale bar (left panel), 50 µm. Scale bar (inset, right panel), 10 µm. N=21 organoids from at least 4 independent batches. Representative image, cell line IMR-90. **F.** Sagittal view of RPE imaged by scanning EM reveals tightly packed single layer RPE cells (yellow, pseudocolored). Primary cilium projects from each cell (red, pseudocolored). Scale bar, 20 µm (left image) and 2 µm (right image). Representative image, cell line IMR-90. **G.** Transmission EM of an RPE cell with numerous melanosomes (yellow arrows) and a primary cilium (white square) **(i).** Magnified primary cilium with a cross-sectioned daughter centriole **(ii).** At least twenty RPE cells were imaged from two independent batches of organoids. Scale bar, Ei, 2 µm and Eii, 1 µm. N=21 organoids from at least 4 independent batches. Representative image, cell line IMR-90. **H-I.** Serial section of developing photoreceptor cilium **(Hi-ii).** Cilium is emerging from a centriole bulging out in the distal area to form an outer segment (OS) suggestive of a rod cell. The OS is distinguishable from the connecting cilium (CC) at the base. Mitochondria is marked MC. An ectosome (red arrow and inset) is released at the tip of the outer segment that takes part in the visual cycle and photoreceptor disc retention and shedding. Right next to the rod cell is an RPE cell with melanosomes. Serial cross-section of a developing photoreceptor with a triangular-shaped outer segment suggestive of a cone cell **(Ii-ii)**. The OS bulging out from the base of the cilium contains several vesicles. An RPE cell is neighboring the cone cell characterized by melanosomes in the cytoplasm. Scale bars, 500 µm. At least nine photoreceptor cells were imaged. N=4 organoids 2 independent batches of organoids. Representative images, cell line IMR-90.

In this region, we could also visualize the layered structure with BRN3-positive RGCs, few α-PKC-positive bipolar cells, PCP4-positive amacrine cells, and calbindin-positive cells, which mark subpopulations of bipolar, amacrine, and ganglion cells in the mammalian retina **(Figure 4C-D and S8B)**. Additional immunostaining using opsin and recoverin antibodies, which specify photoreceptor cells, identified the infrequent presence of positively stained cells **(Figure S8C-D).** Staining for rhodopsin revealed only a diffused staining and never showed a distinct structure (shown later). These data suggest that OVB-Organoids contain cells with identity markers committed to photoreceptor development. Corroborating to this is our scRNA sequencing data revealing the cell clusters expressing outer limiting membrane (OLM) markers (For example, CRB2 and ITGB8). Besides, OLM, CRB2 is localized at the developing photoreceptors(Quinn et al., 2018) **(Figure S2).** We also identified densely packed elongated nuclei from basal to the apical surface facing the outer RPE. This arrangement is not identical but somewhat similar to the inner and outer nuclear cell layers observed in mouse or human fetal retina(Hoshino et al., 2017). Thus, unlike *in vivo* counterpart, OVB-Organoids did not show distinct layers suggesting that they are early-stage optic vesicles containing different cell types but not segregated **(Figure S8E-F).** We observed a positive region marked by cone-rod homeobox protein (CRX), a photoreceptor determinant (Furukawa et al., 1997b). This region was also immunoreactive for L/M-opsin, which specify developing cones. In this region, staining for rhodopsin did provide a vague signal and did not segregate from opsin staining, indicating that it is a developing retinal region (data not shown). Thus, the architecture of OVB-Organoids’ developing neural retina is similar to the early fetal retina (Sridhar et al., 2020). Overall, compared to Capowski et al., stage 3 isolated retinal organoids (Capowski et al., 2019), which display well-defined several outer nuclear layers with segregated rods and cones, optic vesicles of OVB-Organoids constitute only a few outermost layers suggesting that they are still in the process of development. Thus, the described optic vesicles of OVB-Organoids could fit between stages 1 and 2 of retinogenesis.

The isolated retinal organoids differentiated from pluripotent cells do not form genuine optic vesicles, cups, or accurately layered RPEs. This could be because the retinal organoids lack forebrain that is a non-retinal tissue type yet developmentally related to retinogenesis(Capowski et al., 2019). As optic vesicles in OVB-Organoids are attached to the forebrain, we tested an ordered arrangement of RPEs. To image the architecture of RPE, we stained for F-actin and Arl13b, which specify the outer cell membrane and primary cilia. We noticed highly organized pigmented cells displaying the typical ‘honeycomb-like’ morphology, each containing a primary cilium **(Figure 4E).** Tissue clearing and whole-mount imaging followed by 3D reconstruction revealed RPE’s organization in an optic vesicle that is spatially restricted to one pole of the organoid **(Figure S8G and Movie 1)**. Finally, scanning electron microscopy (EM) of transverse sections identified that RPE presents as a tightly packed monolayer with each cell harboring a primary cilium in its surface, an arrangement that is reminiscent of the *in vivo* counterpart **(Figure 4F)**. Disassociation of the pigmented region and subsequent plating also resulted in a monolayer sheet of RPE cells displaying ‘honeycomb-like’ morphology **(Figure S8H).** Basal to pigment-rich RPE, we noticed several serially organized cilia in the rhodopsin-diffused area, suggesting that these Arl13b-positive structures are possibly connecting cilia of developing photoreceptor cells **(Fig. S8I).**

Visualizing these structures using serial sectioning transmission EM revealed distinctive features of RPE cells with numerous cytoplasmic melanosomes(Strauss, 2005). Each RPE cell harbored a primary cilium on its surface **(Figure 4G)**. The inner layer of RPE contained rod-like cells, which displayed a typical bulgy outer segment enclosed by an RPE cell at the apical side and connected to an inner segment via a connecting cilium **(Figure 4Hi and S9A-B)**. Importantly, we also identified ectosomes release from the outer segment, a cellular process that also occurs during outer segment formation(Salinas et al., 2017) **(Figure 4Hii)**. Finally, serial sectioning analysis of a cell revealed different developmental stages of a photoreceptor cilia starting from distal centriole to developing outer segment **(Figure S10).**

In comparing with the developmental stages of ciliogenesis during photoreceptor development of mouse retina, the observed structures appear to fit between stage S4 and S5 of photoreceptor cilia development at which proximal cilium becomes the connecting cilium, and the distal part starts to bulge with membrane accumulation(Sedmak and Wolfrum, 2011). We also noticed another kind of cell type that was morphologically distinct from rods. These cells contained a triangular-shaped outer segment containing many vesicles, flattened cisternae, and mitochondria at the vicinities of their basal bodies, suggesting that they are cone cells. In some cases, we also noticed mitochondria at their base **(Figure 4I and S9C).** In summary, OVB-Organoids contain diverse retinal cell types organized as an immature neural retina.

### OVB-Organoids contain lens- and corneal epithelium-like structures

Based on our scRNA and bulk RNA transcriptomic data, we wondered OVB-Organoids exhibit non-neuronal cell types of lens and corneal epithelium, which are derived from surface ectoderm. Organoids showed two defined bilateral structures strongly positive for αA/αB-Crystallin positioned within each optic vesicle **(Figure 5A).** Importantly, these lens structures are enclosed within a single layer of F-actin and keratin-3-positive columnar epithelial cells suggesting that corneal epithelium neighbors the lens-like structures **(Figure 5B)**. Ultrastructurally, the lens is displayed as a distinctly rounded structure within a space, possibly the anterior chamber enclosed by the corneal epithelium **(Figure 5C and D)**. Lens structures are further characterized by a substantial reduction of cytoplasmic organelles, a process that occurs during the differentiation of lens fiber cells(McAvoy et al., 1999).

**Figure 5.**
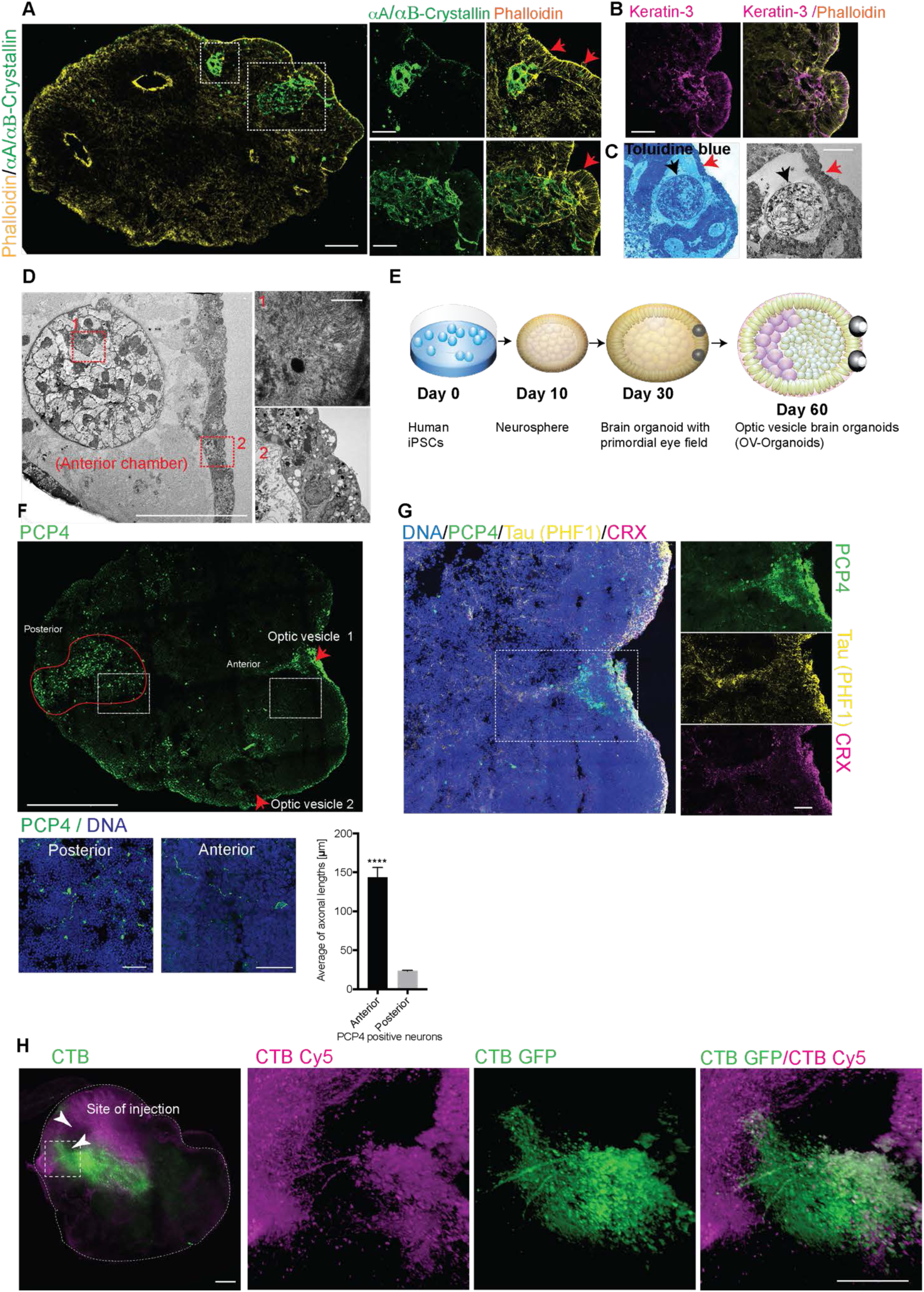
OBV-Organoids brain organoids display non-neuronal components and optic tracts connected to brain organoids. **A.** A 60-day-old OVB-Organoid stained with F-actin marker Phalloidin (yellow and arrows). Two lenses are stained by a lens marker *α*A/*α*B Crystallin (green) Scale bar, 100 µm. Magnified images are at right. Scale bar, 50 µm. Representative images are shown. N=10 organoids from at least 3 independent batches. Representative images, cell line IMR-90. **B.** Lenses (green) are neighbored by a single epithelial cell layer that is Keratin-3 positive (magenta). Representative images are shown. Scale bar, 50 µm. N=7 organoids from at least 3 independent batches. Representative images, cell line IMR-90. **C.** Ultrathin section of the optic vesicle region stained with toluidine blue staining showing a well-defined round lens (black arrow) in a cavity. Red arrow points at the cell monolayer of the cornea on the surface next to the lens. Panel at right is an EM image of the similar region with lens (black arrow) and cornea (red arrow). Scale bar, 50 µm. Representative image, cell line GM25256. **D.** Magnified view of the same lens. Anterior chamber enclosed by the cornea. Square 1 shows an area where the transition from the lens vesicle to lens fibers happened. The lens fibers are visible in inset 1 (upper right panel). Inset 2 (lower right panel) shows magnified corneal cell layer with typical tight junctions. Cornea is separated by a basal membrane (probably early Descemet’s membrane) from the inner chamber. Scale bar left image, 100 µm. Scale bar insets, 10 µm. **E.** Schematic of OVB-Organoids generation **F-G. (i)** Whole organoid section stained for PCP4 (green), which is expressed in the retina, developing lateral geniculate nucleus (LGN) and visual cortex (area 17). PCP4 signals (green) in the optic vesicle region (red arrow, optic vesicle 1), whose axons merge to a possible optic stalk (the precursor of the optic nerve), and in cells of the posterior region of the organoid (red circled area). Scale bar, 500 µm. **ii.** Right images show magnified optic vesicle 1 area with CRX (magenta) and PCP4-positive retinal region (green). Bar, 100 µm for merged image and 50 µm for insets. Magnified panels at right show PCP4 region merges with optic stalk where axons are PHF1 (adult form of Tau) positive (yellow) **(G)**. Scale bar, 100 µm for merged image and 50 µm for insets. Panels at right shows magnification of squares in the anterior and posterior region of the organoid with neurons positive for PCP4 (green). Scale bar, 50 µm. Panels below shows quantification of axonal lengths of PCP4 positive neurons in the anterior and posterior region revealed that anterior PCP4 positive neurons possess significantly longer axons than those of the posterior region. Axons of 100 neurons from 3 different batches of organoids were measured. Unpaired t-test One-way ANOVA, p****<0.0001, n = 3. Error bars show +/- SEM. Representative image, cell line IMR-90. **H.** CTB injection experiment. Lower panel: Representative volume-rendered images of organoids injected with Cholera Toxin B (CTB-488 and CTB-647) at two different optic vesicles of the same organoid. White arrowheads mark the site of injection. Reconstructed images of individual channels, as well as merged channels, are shown at right. Scale bars, 200 µm. N=13 organoids from 4 independent batches. Cell line IMR-90.

Indeed, the lens cells contained cytoplasmic aggregates of fibrillar material **(Figure 5D).** Intriguingly, these observations are reminiscent of *in vivo* tissues observed at stage 23 of prenatal development (O’Rahilly, 1966, 1975). In summary, with the application of low seeding cell density in a defined but undirected differentiation condition, we could engineer brain organoids with bilaterally symmetric optic vesicles. Although the described *in vitro*-tissues do not fully mimic *in vivo*-like architecture, they display diverse cell types of complex prenatal eye development **(Figure 5E)**.

### OVB-Organoids contain primitive image forming nuclei and exhibit retinal connectivity to brain regions

In the mammalian brain, axons of retinal ganglion cells reach out to connect with their brain targets, an aspect that has never been shown in an *in vitro* system. First, we tested for the lateral geniculate nucleus (LGN) presence, which is the primary image forming nuclei of the visual thalamus that develops when eye-specific axonal segregation is completed (Sretavan and Shatz, 1986). We identified immunoreactivity for PCP4, a developmental marker of LGN, amacrine, and bipolar cells at the vicinities surrounding the optic vesicle suggesting that PCP4 could also specify ganglionic cells (Chintalapudi et al., 2016) **(Figure 5F)**. PCP4-positive cells showed a typical morphology of cell soma with an axon extended up to 100µm. The PCP4-positive structures penetrated way out of the optic vesicle migrating from anterior to posterior of the organoid. The distinct PCP4 clusters posterior to the optic vesicle suggest that PCP4-positive neurons could have migrated from the retina to form higher-order visual regions that are yet to be characterized. Intriguingly, staining for Tau, an axonal marker, also co-localized with the extending PCP-4 fibers migrating towards the inner region of organoids. Thus, the regions demarcated by PCP4- and Tau-is, to some extent, morphologically similar to the primitive optic disc and optic stalk structures **(Figure 5G).**

To further test that optic vesicles contain projection tracts into organoid tissues, we microinjected AlexaFluor488-cholera toxin b-subunit (CTB) directly into one optic vesicle and AlexaFluor647-CTB into the other optic vesicle. CTB labels retinal nerve fiber of axon bundles when injected to eyes (Huberman et al., 2005) (Angelucci et al., 1996) (Crish et al., 2010). After 24 hours of light exposure (see methods for details), we noticed a strong uptake of CTB into the retinal region with numerous axon-like projections **(Figure 5H and S11).** 3D reconstruction of confocal slices revealed that the fiber-like optic tracts merge, although they appear to emerge from two different optic vesicles **(Movie 2).** These observations align with our transcriptomic analysis and raise the possibility that the optic vesicles are connected to brain organoids’ inner regions via axonal-like projections.

### OVB-Organoids show signatures of cortical neuronal maturation and functionality

From our transcriptomic data, we inferred that OVB-Organoids strongly correlate with the development of a neural retina in the optic vesicles of OVB-Organoids, while also exhibiting signatures of synapse maturation and myelination of cortical neurons **(Figure S5)**. Testing this, indeed, OVB-Organoids strongly expressed Synapsin1-positive region restricted to cortical plates and is also specified by mature neuronal markers like CTIP, Myelin Basic Protein (MBP), and laminin **(Figure 6A-D).** Consistently, our GSEA data revealed a down-regulation of genes regulating cell proliferation, indicating the quantitative commitment of these organoids to maturation. Corroborating these findings, OVB-Organoids exhibited an enrichment of cells expressing genes for sensory perception of light stimulus, strengthening the presence of mature functional cell types **(Figure 6E-J).** To substantiate the existence functional neurons, we performed whole-cell patch-clamp recordings. In on-cell configuration, spontaneous action potentials could be measured, which were sometimes sensitive **(Figure 6K)** and occasionally resistant to Tetrodotoxin (TTX). TTX is a neurotoxin that selectively blocks sodium channels (Makarova et al., 2019). The resistance to TTX treatment is typical for retinal cells, which possess TTX-resistant voltage-gated sodium channels (O’Brien et al., 2008).

**Figure 6.**
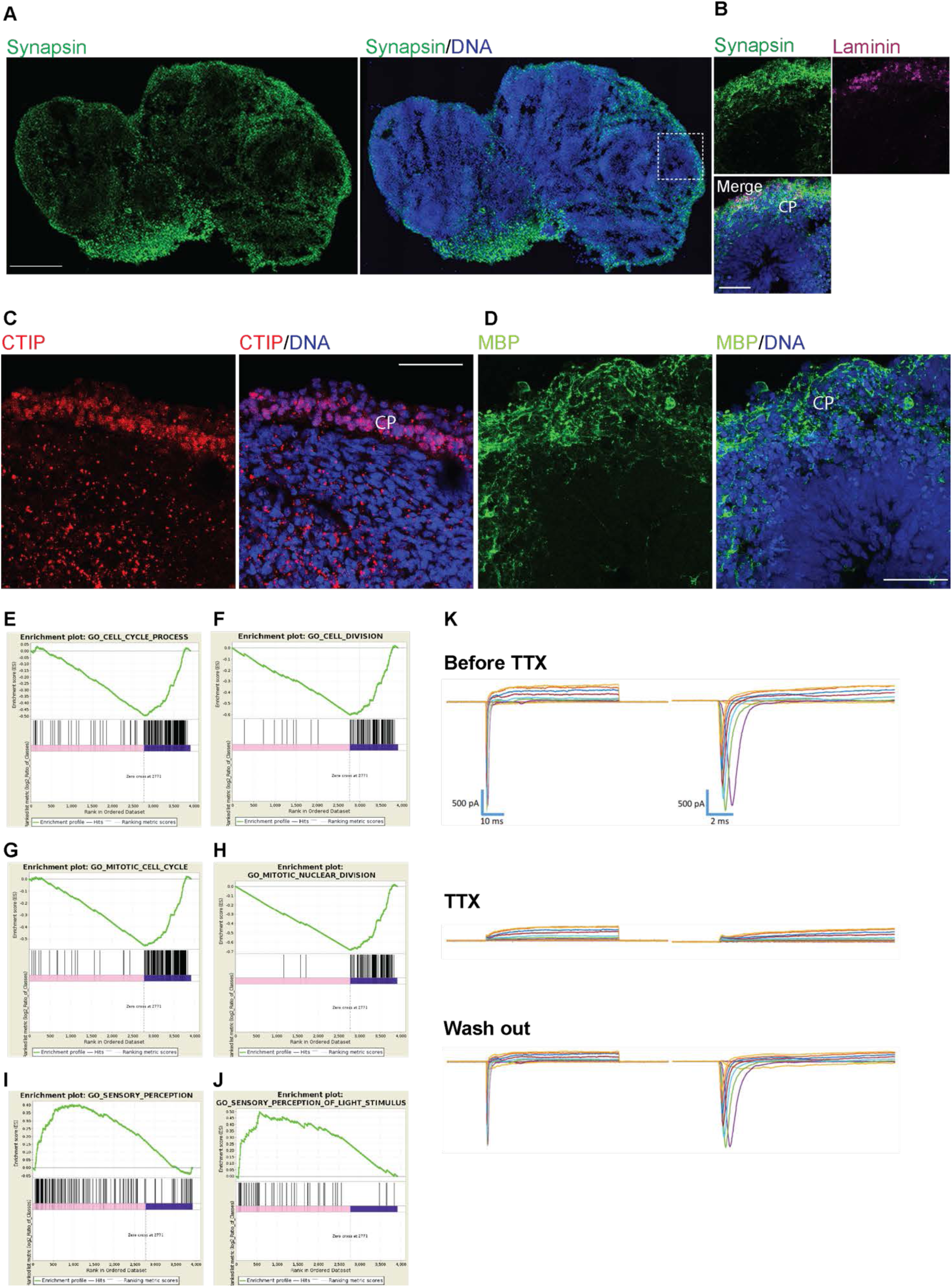
OBV-Organoids display mature neurons. **A-B.** Section of the whole organoid stained for Synapsin (green) and Laminin (magenta), proteins essential for the release of synaptic vesicles at the presynaptic terminals, and polarization and layering of the neurons in the cortical region, respectively. CP, cortical plate. Scale bar, 500 µm (overview of the whole organoid) and 50 µm (inset). Representative images are shown. N=17 organoids from at least 5 independent batches. Representative images, cell line IMR-90. **C-D.** Cortical region of a brain organoid exhibits a CTIP2 (neurons of layer V) positive cells layer (red). Scale bar, 25 µm. MBP (myelin basic protein, green) positive region indicating the presence of myelin-producing oligodendrocytes **(D).** Representative images are shown. N=12 organoids from at least four independent batches. Representative images, cell line IMR-90. **E.** Enrichment plot showing cell cycle processes down-regulated in OVB-Organoids, indicating higher numbers of matured cell types compared to early brain organoids. Cell line IMR-90. **F.** Enrichment plot showing cell cycle division is down regulated in OVB-Organoids indicative of a higher number of post-mitotic cells compared to early brain organoids. **G.** Enrichment plot showing mitosis genes are down regulated, as shown in panels E-F. **H.** Enrichment plot showing OVB-Organoids contain cells that mostly have down-regulated the genes for mitotic nuclear division in contrast to fast-cycling progenitor cells in early brain organoids. **I.** Enrichment plot showing OBV-Organoids contain cells with up-regulated gene expression for sensory perception. **J.** Enrichment plot showing OVB-Organoids contain cells expressing genes for sensory perception of light stimuli. Results from panels **(I)** and **(J)** support the presence of matured functional cell types relevant to eye development in the OVB-Organoids compared to early brain organoids. **K.** Two currents (inward and outward) are seen (about 400pA at +40mV). The outward current is TTX insensitive and is an outwardly rectifying K+ current. Whole-cell patch-clamp recording revealed that neurons show a transient inward TTX (Tetrodotoxin) sensitive current. TTX is a neurotoxin that selectively blocks the sodium channel. Here is shown one recording example of a neuron with a high Na+ current. Upper two graphs show the recorded current before treatment. Middle graphs show blocked Na+ current (inward current) by TTX treatment. And bottom panels show reoccurring Na+ current after washing out of TTX. Scaling bar is 500 pA. Total recorded cells were 27 and 36 per organoid from two independent batches. Cell lines IMR-90 and GM25256.

### OVB-Organoids are light sensitive and can recover their light sensitivity after photobleach blinding

To test the functionality of OVB-Organoids, we performed electroretinography (ERG) recordings that allow the quantification of retinal signaling. In the absence of light, vertebrate photoreceptors are slightly depolarized, and light absorption leads to membrane hyperpolarization (Schiller et al., 1986). (Yamaguchi et al., 1992) Typically, the first graded electric response represents the photoreceptors’ hyperpolarization and is quantified as the a-wave response. Trans-synaptic excitation spreading mainly leads to the depolarization of ‘ON’-bipolar cells, modulated by the neuronal network between photoreceptors and ganglion cells. The resulting net depolarization is recorded as the ERG b-wave.

We recorded a response to every 500ms white light flash at an interval of 3 min (see method section for details). To quantify neuronal signaling, we calculated the amplitudes and their implicit time. Notably, a dose-dependent light exposure already revealed increasing negative amplitudes of the ERG corresponding to increasing light intensities **(Figure 7A).** The first question that arose in this experiment was: Does the negative deflection represent photoreceptors’ activity solely, or are there apparent negative waves containing a positive b-wave deflection? Indeed, in the vertebrate retina, the positive b-wave responses superimpose the negative a-wave response (Albanna et al., 2009) (Yamaguchi et al., 1992).

**Figure 7.**
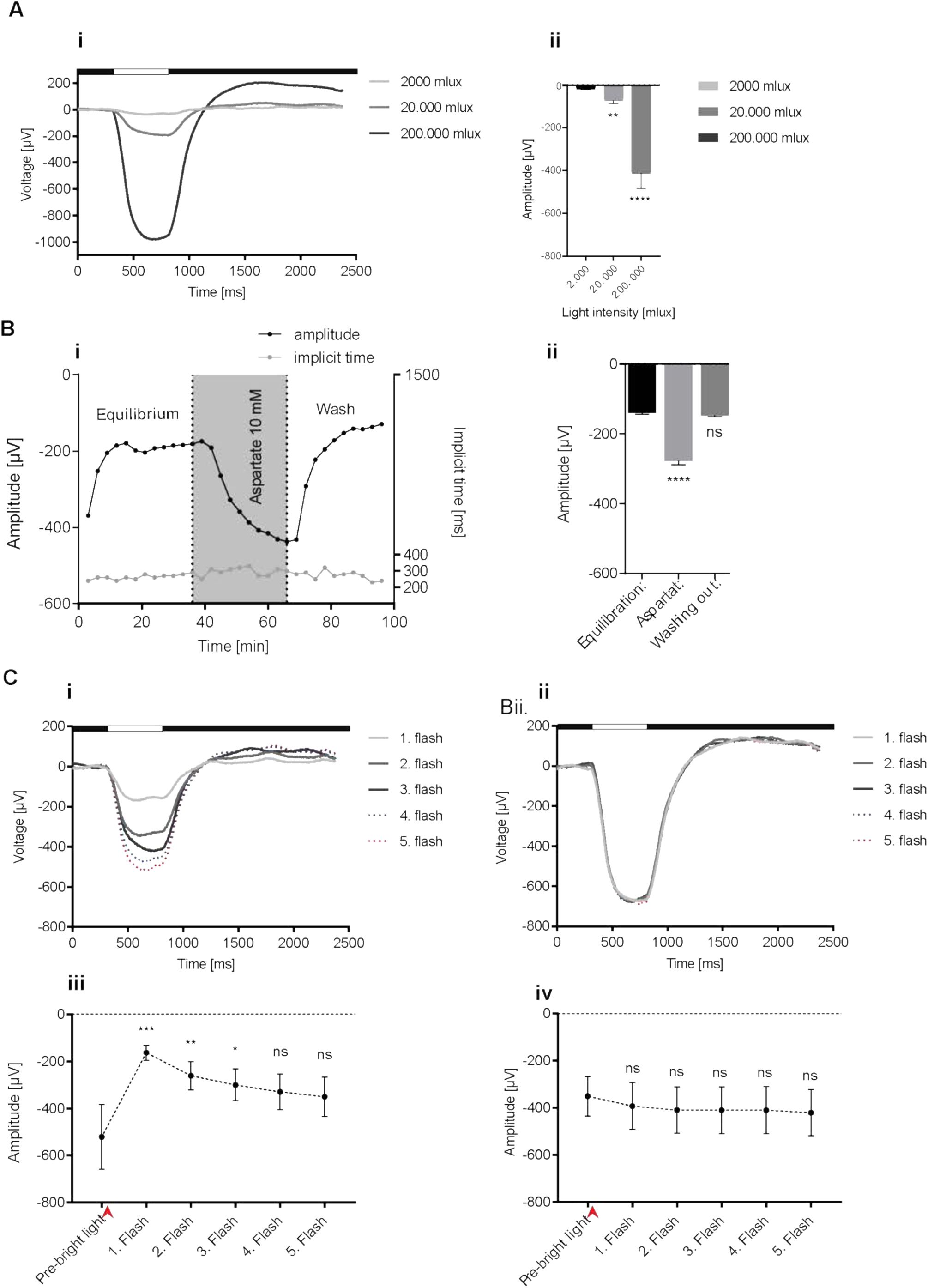
OVB-Organoids are light sensitive. **A. (i).** A dose-dependent light response stimulated with three different light intensities. The white line at the top of the graph marks the time span of light flash (500 ms). **ii.** Statistical summary. Each organoid was exposed and recorded every 3 minutes for 3x 500 ms to 2000 mlux (∼candle light), then 3x 500 ms to 20.000 mlux (∼street light at night) and finally 3x 500 ms to 200.000 mlux (∼sunset or sunrise on a clear day). Light response of organoids increased significantly with increasing light intensity (20.000 mlux compared to 2.000 mlux p<0.001, 200.000 mlux compared to 2.000 mlux p<0.00001). Kruskal-Wallis test of One-way ANOVA data: non-parametric; Dunńs multiple comparisons test; mean with error bars SEM; n=9. Cell line IMR-90. **B.** Isolating A-wave within the retinal signaling network. **i.** ERG upon aspartate treatment. During equilibration time of 36 min, organoid got adjusted to the system under 2 ml/min superfusion and the light flash of 500 ms with 200.000 mlux light intensity until it showed the stable amplitude of −200 µV. Within 6 min of aspartate application OVB-Organoid showed a drop in amplitude up to lesser than - 400 µV upon light flashes. After aspartate washout, organoid shows amplitude that is similar to aspartate-free condition. **ii.** Statistical summary of aspartate treatment. 10 mM aspartate treatment led to a significant hyperpolarization of −278 µV on average compared to equilibration before aspartate treatment (−140 µV on average) (****p<0.0001). Hyperpolarization after washout of aspartate is similar to value before aspartate application (−147 µV). Error bars show mean +/- SEM. One-way ANOVA data followed by Dunnett test. N = 3 organoids. Cell line GM25256. **C.** Photosensitivity of OVB-Organoids can be desensitized by bright light exposure with the light intensity of 4600 Lux for 10 minutes **(i)**. White line at the top of the graph marks the time span of light flash (500 ms). **(ii).** Negative control experiment. The same organoid was treated after 30 min recovery from photic stress under the same conditions in which bright light exposure was replaced by 0 lux. **iii.** Statistical summary of photic stress experiments. OVB-Organoids in electrode chamber were adapted for 15 min to the ERG system (Pre-bright light) with light flashes (200.000 mlux) and recording every 3 minutes. Then organoids were photically stressed by exposure to 4600 lux for 10 minutes (red arrow). While average amplitude before bright light exposure was - 521 µV, photic stress led to a significant reduction of photosensitivity (p<0.0001 after 30 s of recovery, p<0.001 after 3:30 min of recovery and p<0.01 after 6:30 min of recovery). Finally, there was no significant difference in light response detectable after 9:30 min and 12:30 min of regeneration suggesting a complete recovery from photic stress within this time period. Friedman test of One-way ANOVA data: non-parametric; Uncorrected Dunńs test; mean with error bars SEM; n = 4. **iv.** Negative control experiment. After recovery from photic stress, each organoid was exposed to the same treatment in which the bright light was replaced by 0 lux (red arrow). No significant difference was detected in photosensitivity, suggesting that the response detected in graph C was solely due to bright light exposure. Friedman test of One-way ANOVA data: non-parametric; Uncorrected Dunńs test; mean with error bars SEM; n = 4. Organoids were superfused in organoid medium with a speed of 0.5 ml/min.

Therefore, to measure photoreceptors’ response, we inhibited trans-synaptic signaling using a glutamate receptor antagonist to detect the full-length amplitude of the a-wave. ERGs in the presence of 10mM aspartate blocked the b-wave, which eventually increased the negative amplitude of the fully developing a-wave to its maximum value. However, during the consecutive washout of aspartate, we could observe the re-occurrence of positive b-wave, indicating the presence of trans-synaptic signal transduction in organoids **(Figure 7B).** This finding demonstrates that the changes in the electric potential represent mainly the responses of photoreceptors but only when aspartate is present. Removing aspartate revealed the positively deflecting b-wave, indicating that trans-synaptic signaling is present in the organoids.

Photobleaching by an intense flashlight stimulation can inactivate physiologically active photoreceptors, which can spontaneously be reversed to their initial responses by dark adaptation (Ernst and Kemp, 1979). To investigate whether organoids reflected this physiological phenomenon, we photic-stressed them by exposure to an increased light intensity of 4600 lux for 10 minutes (“Pre-bright light”). The amplitude of the electrical responses was transiently reduced after light stressing when recorded with short, low-intensity recording light pulses. Interestingly, the organoids could recover their photosensitivity as we could detect normalized electric responses during consecutive dark adaptation **(Figure 7C)**. These experiments reveal that organoids contain physiologically active photoreceptors that are capable of recovering light sensitivity after photobleaching. Importantly, when repeating the photic stress experiment with isolated mouse retina lacking pigmented epithelium, this kind of recovery of the photosensitivity was not observed **(Figure S12).** This finding highlights the importance of complexity and interactions between different cell types present in optic vesicles that are functionally integrated within the organoid. In summary, most aspects of our recording experiments strongly correlate with our transcriptomic and imaging experiments.

## Discussion

We aimed to engineer a complex interorgan *in vitro* system such as human brain organoids capable of assembling functionally integrated optic vesicles. Here, we showed that iPSC-derived brain organoids could assemble optic vesicles and generate various neuronal, non-neuronal, and retinal compartments within a single organoid. Due to the assumption that brain organoids are not able to strictly follow self-patterning rules like embryos, brain organoids are instead thought to remain as chaotic 3D tissues lacking anterior-posterior and dorsal-ventral axes. Recent elegant work identified that organoids could be induced with gradients of signaling molecules to define spatial topography of the forebrain (Cederquist et al., 2019). Here, we demonstrate that organoids can spontaneously develop bilaterally symmetric optic vesicles from the forebrain-like region without artificial signaling centers.

While the mechanisms that govern spatial topography to establish bilaterally symmetric optic vesicles require future investigations, the presented work already highlights the intrinsic self-patterning ability of iPSCs in a highly complex biological process. Nonetheless, the current work offers at least two indications as an indirect consequence of bilateral symmetric optic vesicles. First, our single sequencing data reveals that the subclusters SC4 to SC6 express ZIC2 and SLIT2 transcription factors determine the routing of RGC axons at the optic chiasm to respective hemispheres, a patterning event that is critical for binocular vision **(Figure 3G)** (Lee et al., 2008) (Herrera et al., 2003). Second, we showed the presence of both CTB-647 and CTB-548-labeled projections when these tracers were injected into two different optic vesicles **(Figure 5H).**

Earlier works by the Sasai laboratory have pioneered optic cups from human embryonic stem cells, which displayed stratified neural retina after >260 days of culturing (Nakano et al., 2012). Later works by Zhong et al. and several other groups have generated similar retinal organoids and demonstrated that they are light-sensitive (Cowan et al., 2020; Zhong et al., 2014). Further studies have reported that the generation of similar retinal organoids is versatile across several independent pluripotent cell lines (Capowski et al., 2019). These works have focused on generating pure retinae. Thus, the functional integration of these 3D retinal structures into brain organoid tissues has remained untested. A critical effort has been made culturing brain organoids for nine months. Even though these organoids did not display polarization, stratification, or visibly apparent RPE rich distinct optic vesicles or vesicles, the differentiation condition led to the generation of photosensitive cell types (Quadrato et al., 2017).

The brain organoids described here display both neuronal and surface ectodermal sublineages. It is unexpected to observe lens and cornea when iPSCs are directed to differentiate into neuroectoderm. Our sc-sequencing analysis potentially addresses the identity and the origin of these non-neuronal cell types. The cell clusters harboring signatures of early eye development (C5 and C11) also contained signatures of preplacodal ectoderm (PPE) and lens ectoderm cell clusters (DLX5, 6, BMP7, MAB21L1, MEIS1 and 2) **(Figure S2).** Thus, the lens and corneal cell types could have originated from the PPE of OBV-Organoids since the PPE contains both neurogenic and non-neurogenic placodes, which give rise to ocular surface ectoderm (Lleras-Forero and Streit, 2012). Second, we did not use retinoic acid (RA) before the neuroectoderm expands, and thus we tried not to force the development of purely neural cell types at this initial stage (Janesick et al., 2015). We believe the timing and the dosage level of RA could influence the co-development of neural and non-neural ectodermal cell types, which requires further investigation (Rosen and Mahabadi, 2020) (Cvekl and Wang, 2009). Finally, it is maybe noteworthy that the function of PAX6 in the eye and brain development is context-dependent as it is also required for the formation of the ocular lens in the eye field region (Duncan et al., 2004) (Oron-Karni et al., 2008).

Intriguingly, when optic vesicles were observed in an organoid, their cytoarchitectures and functionality were similar regardless of which cell line was used to generate the organoid. Importantly, these organoids’ optic vesicles are functionally integrated as demonstrated by axon-like projection tracts and electrically active neuronal networks. The time frame to generate these complex structures (from 50 to 60 days) is a crucial step forward *in vitro* developmental neurobiology as it fulfills the practical feasibility of using this system in multiple experimental setups in a reasonable time window. Notably, the generation of various cell types organized as an immature stratified neural retina within 60 days parallels human embryo (O’Rahilly, 1966, 1975). Table 5 summarizes key differences between different methods. Further refinement of the described method will help reconstruct early brain development complexities in parallel with optic vesicle morphogenesis and its circuitry.

**Table 5:** Summary of key differences between different methods used to generate retinal organoids

In our organoids, we could apply ERGs, a diagnostic test method, to show strong hyperpolarizing a-wave, strictly connected with the early photoreceptor response, after eliminating the b-wave. Furthermore, our photic stress experiments indicating that photoreceptors can be regenerated **(Figure 7).** The measured rapid response within less than 500 ms suggests that the response is a photoreceptor-specific reaction and reveals vertebrate classical photoreceptor cells’ characteristics but not intrinsically photosensitive retinal ganglion cells (ipRGCs), which would depolarize but not hyperpolarize upon light stimulation (Johnson and Pak, 1986). However, the typical light-dependent responses are recorded at an about thousand-fold higher light intensity as found with native isolated bovine or murine retina. This may reflect the relatively low expression density of typical photoreceptor characterizing proteins in these systems or the lack of dark adaptation systems in our organoids, similar as in humans with congenital stationary night blindness (Al Oreany et al., 2016). However, it does not exclude the presence of developing vertebrate photoreceptors rather than ipRGCs. Future experiments that involve genetic ablation of essential phototransduction components and subsequent recordings are required to confirm that retinal photoreceptors purely elicited the observed responses.

OVB-Organoids displaying highly specialized neuronal cell types can be further developed, paving the way to generate personalized organoids and RPE sheets for transplantation therapies. Indeed, we showed that RPE cells could also be cultured from OVB-Organoids **(Figure S8H).** Finally, we believe that OVB-Organoids represent the next-generation organoids helping to model retinopathies that emerge from early neurodevelopmental disorders.

## Acknowledgement

We thank Eugenio F. Fornasiero for the fruitful discussions and support. We want to thank Dieter Häussinger and Boris Görg for offering their support with their microscope facility. We thank E. Paccagnini for the help with the scanning electron microscope. We acknowledge our use of the gene set enrichment analysis, GSEA software, and C5 Molecular Signature Database (MSigDB) (Subramanian, Tamayo, et al. (2005), PNAS 102, 15545-15550, http://www.broad.mit.edu/gsea/). We are thankful for PHF-1 antibody that was kindly provided by Peter Davies. We thank the lab members of the Laboratory for Centrosome & and Cytoskeleton Biology and the Institute of Human Genetics for their support. This study was supported by the Priority Programme „Gene and Cell-Based Therapies to Counteract Neuroretinal Degeneration“(SPP 2127) and by the Fritz Thyssen Stiftung (E.G. and J.G.). This study was supported by the German Research Council (SFB974, RTG1949), and by the Jürgen Manchot Graduate School MOI (P.A.L. and P.V.S.).

## Author contributions

J.G., E.G.: Conception and study design, organoid protocol development, data analysis and interpretation, writing the manuscript, E.G.: hiPSC culture, differentiation, characterization assays, organoid generation, RPE isolation and culture; G.P., V.B.: single cell RNA sequencing analysis; J.A., sc-sequencing; C.M.K for iPSC lines. G.P and V.B.: sc-RNA transcriptomics and analysis; W.A., T.S.: W.A., T.S.: Designing and performing ERGs and analyses, photic stress and aspartate experiment; A.P., N.J.: RNA-seq experiments and analyses; A.R.: Designing and performing microinjection, interpretation and analysis; A.R, A.M.: 3D confocal imaging and 3D movies; G.C., M.R., V.P.: TEM and SEM, analysis; M.G.: TEM and analysis; E.G., F.S.: fluorescent imaging experiments and analysis; G.B., S.O.R.: Designing and performing patch clamping experiments and analysis; J.H., A.R., A.M., proofread the manuscript.

## Supplementary figures and legends

**Fig. S1.**
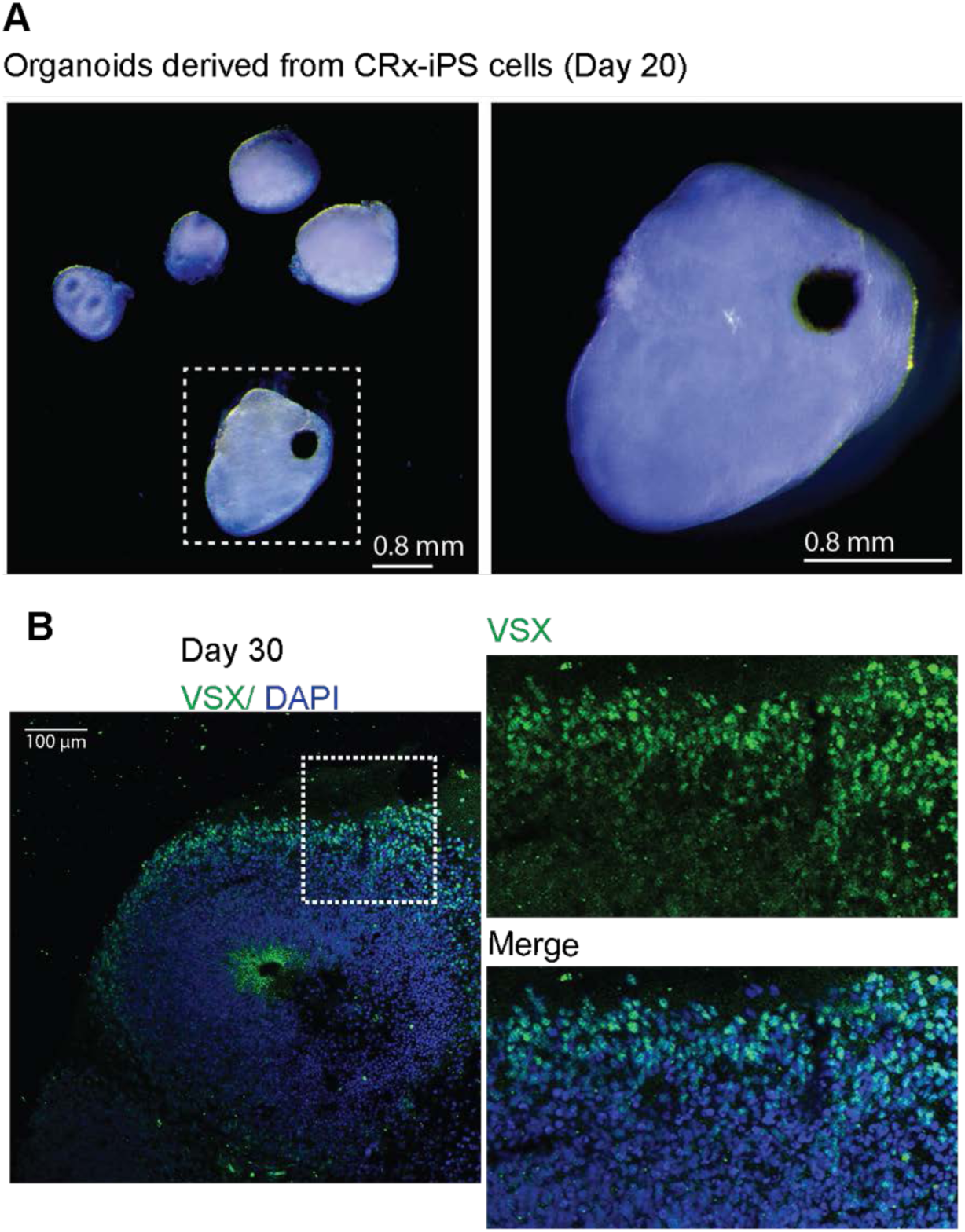
Related to main figure 1. **A.** Organoids differentiated from Crx-iPS lines reprogrammed from adult retinal müller glial cells generates strongly pigmented structures within 20 days. N=18 organoids from at least 3 independent batches. Representative images, cell line Crx-iPS. **B.** Day 30 organoids exhibits immunoreactivity of VSX, progenitors restricted at the primitive eye field. Representative image is given. N=15 organoids from two independent batches. Cell line IMR-90.

**Figure S2.**
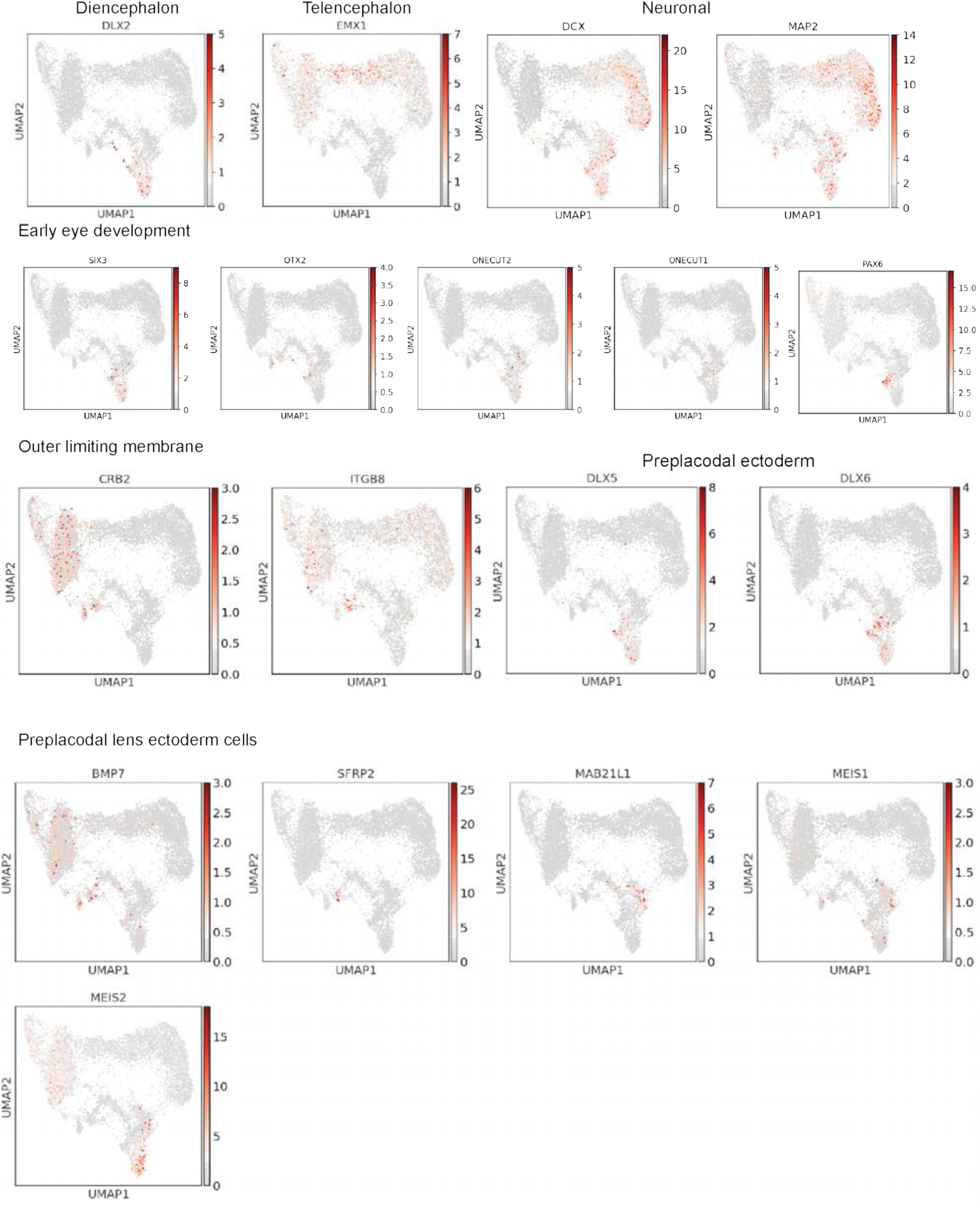
Related to main figure 3. Feature plots show expression of representative genes used to define major cell clusters in 60-day OVB-Organoids. Diencephalon (specified by DLX2), Telenchephalon (specified by EMX1), Eye development (specified by SIX3, OTX2, ONECUT1/2 and PAX6), Outer limiting membrane (specified by CRB2 and ITGB8), Preplacodal ectoderm (specified by DLX5 and 6) and Preplacodal lens ectoderm cells (specified by BMP7, SFRP2, MAB21Li, MEIS1 and MEIS2.

**Figure S3.**
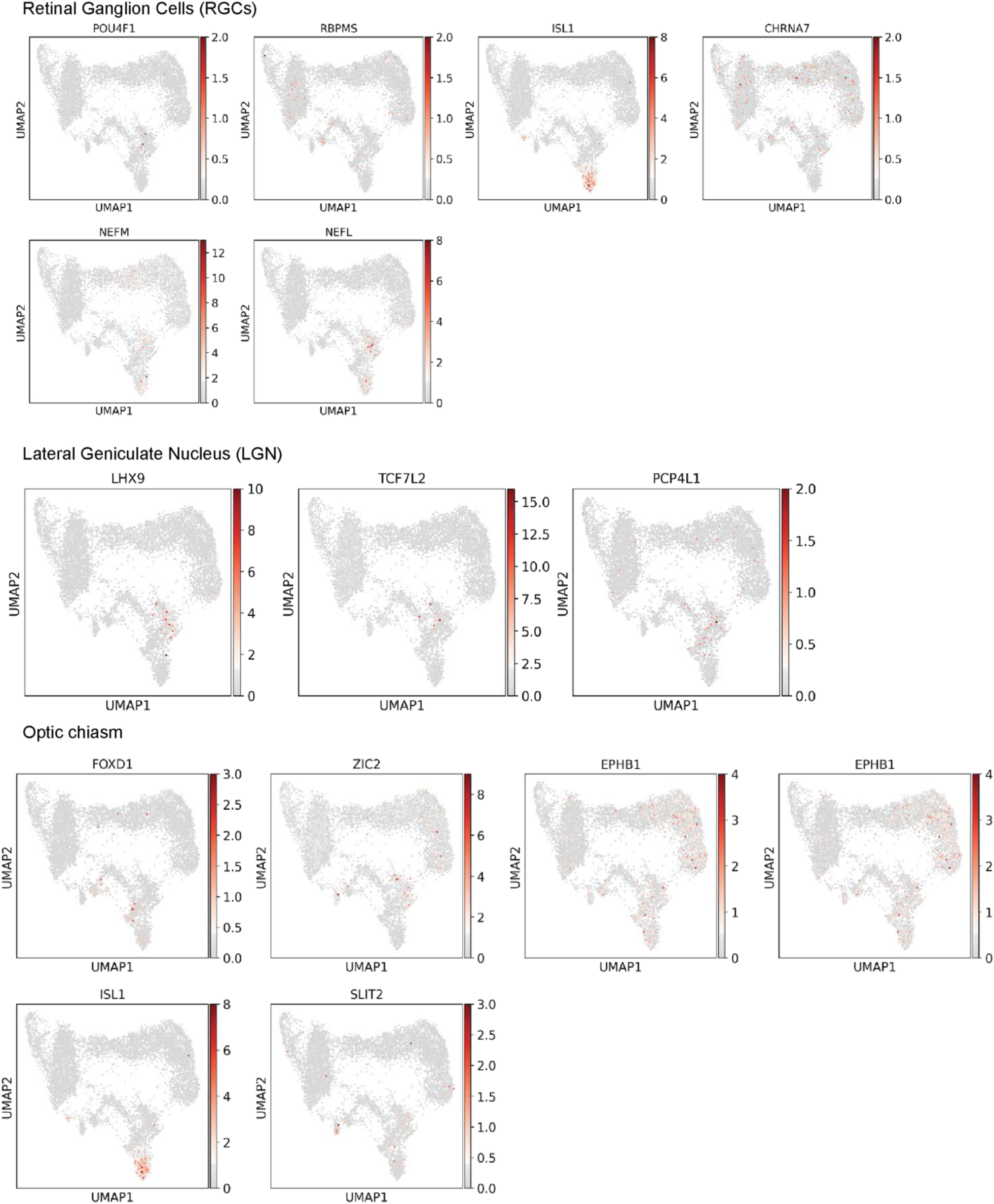
Related to main figure 3. Feature plots show expression of representative genes used to define major cell clusters in 60-day OVB-Organoids. Retinal ganglion cells (RGCs) (specified by POU4F1, RBPMS, ISL1, NEFM and NEFL), Lateral Geniculate Nucleus (LGN) (specified by LHX9, TCF7L2, PCP4L1) and Optic chiasm (specified by FOXD1, ZIC2, EPHB1, ISL1 and SLIT2).

**Figure S4.**
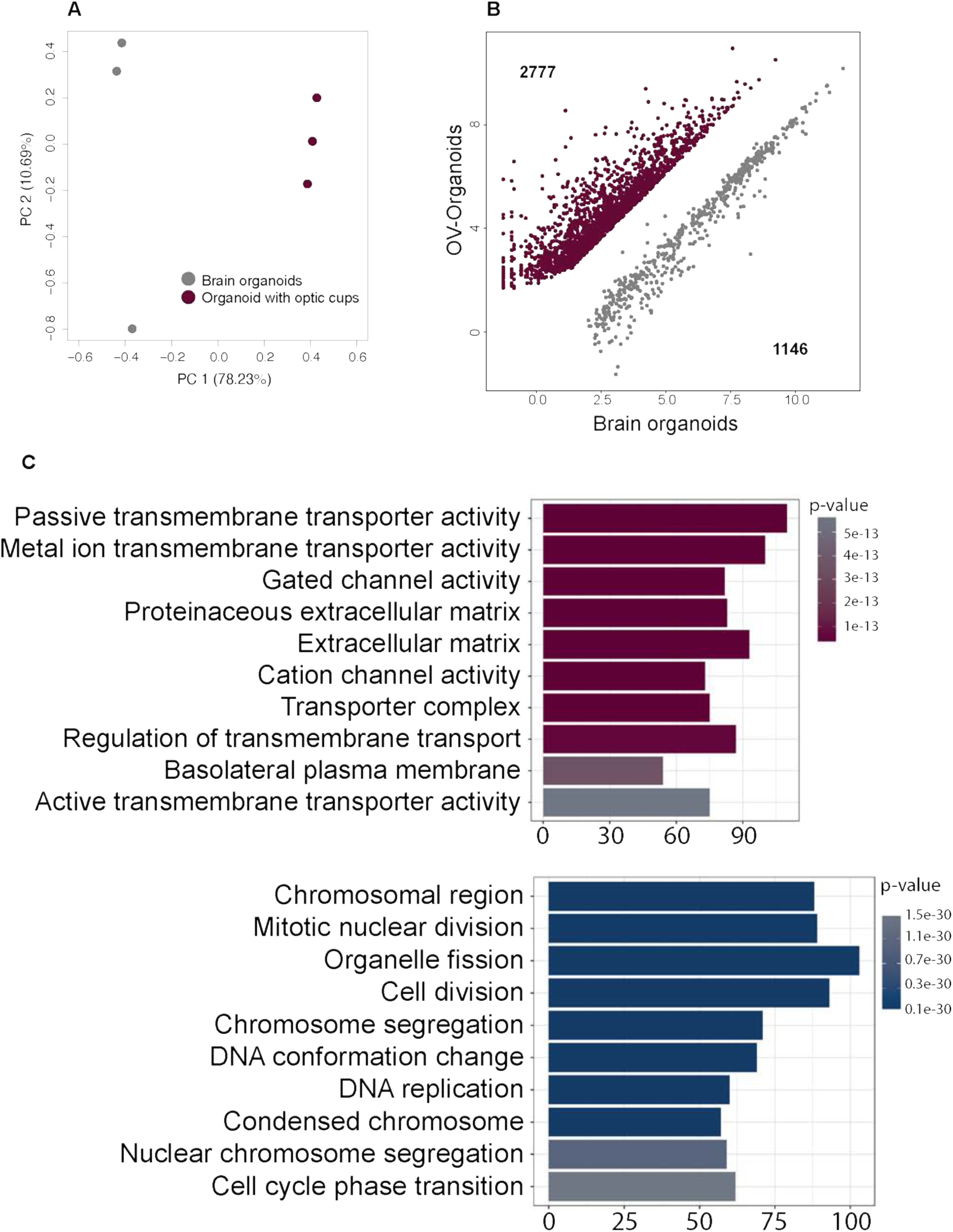
**A.** Analysis of mRNA-seq data from early brain organoids with primitive eye fields and OVB-Organoids (both generated from cell line IMR-90). Principal component analysis biplot of mRNA-seq data derived from brain organoids and OVB-Organoids. PC1 contributes to ∼80% of the variance between the sample subgroups. **B-C.** Scatter plot of normalized counts from mRNA-seq data reveals 2777 genes (*purple*) being up-regulated and 1146 (*grey*) down-regulated in OVB-Organoids compared to early brain organoids with primitive eye fields.

**Figure S5.**
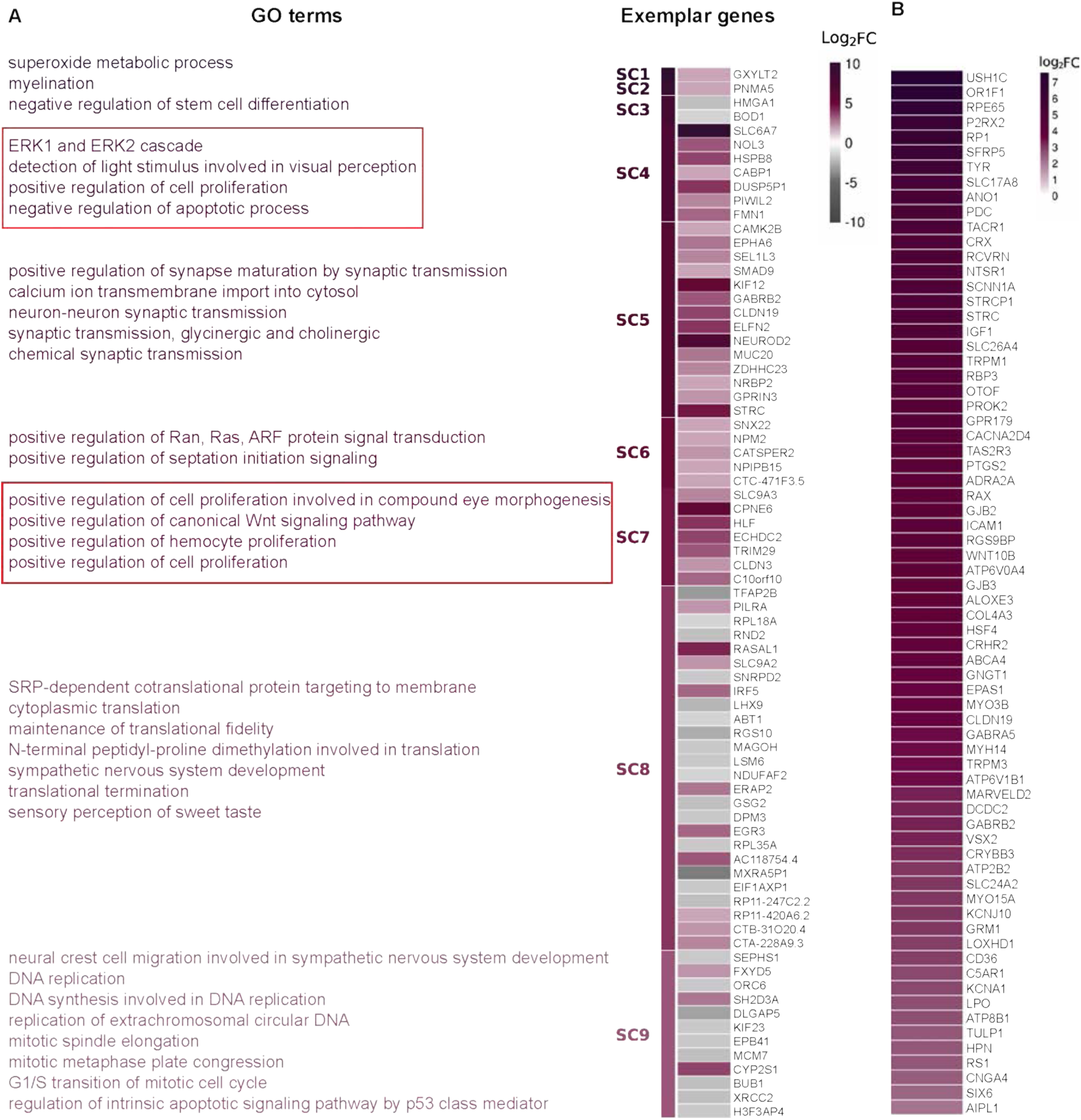
**A.** Affinity propagation clustering results represented by the log_2_ fold-changes of exemplary genes from 9 super-clusters and their respective gene ontology terms indicating super-clusters 4 and 7 as clusters with genes involved in visual perception and eye morphogenesis. **B.** Leading-edge analysis of GSEA results for super-cluster 4 reveals a retinal development gene signature enrichment, exemplified by retinal progenitor and ganglion markers, as well as typical photoreceptor cells transcription factors.

**Figure S6.**
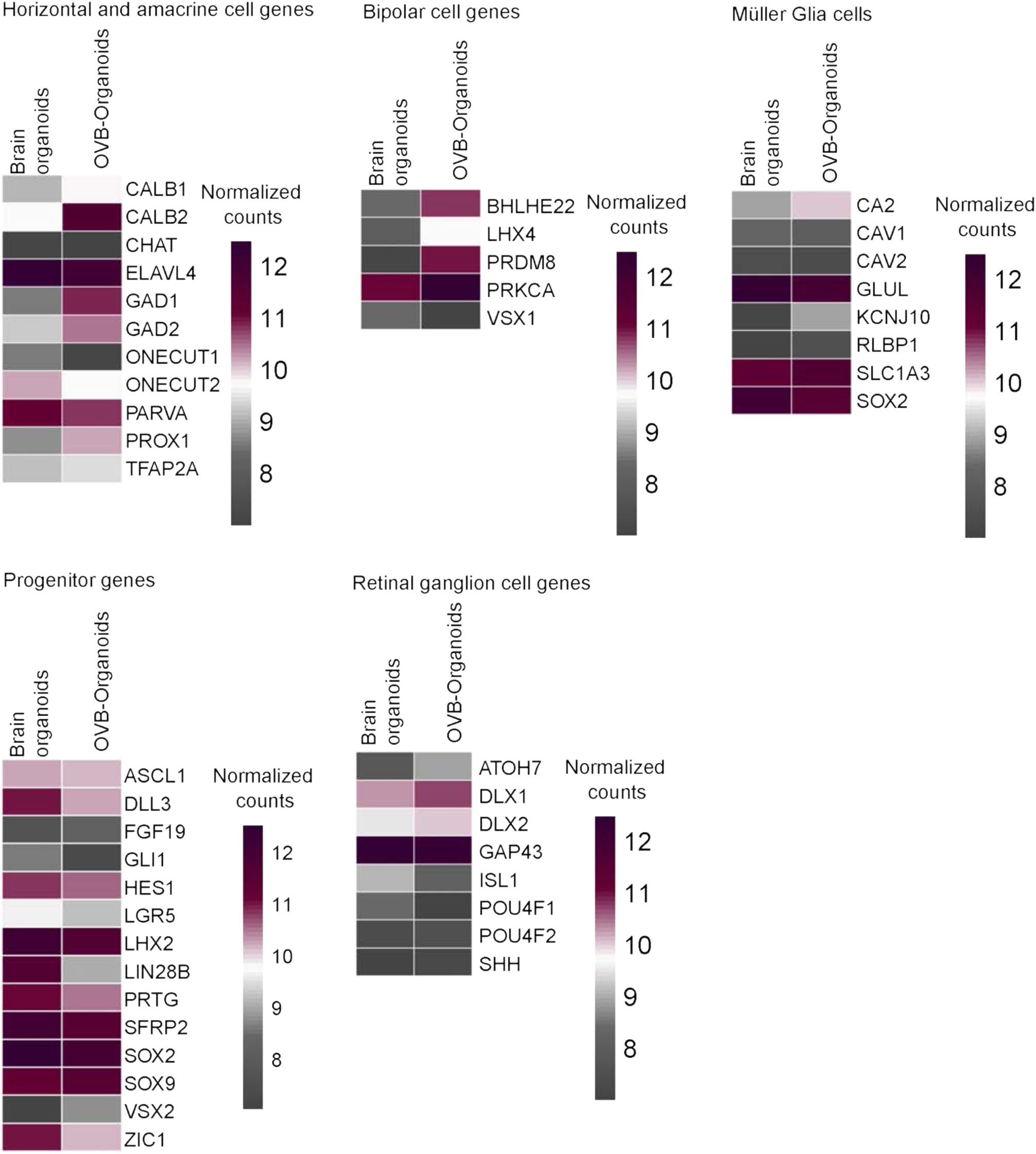
Expression profiling of OVB-Organoids shows developmental eye signatures. The mRNA-seq analysis of early brain organoids with primitive eye fields versus OVB-Organoids is shown compared to human fetal retina transciptome data (STAR aligner Hg19 human reference transcriptome) Genes are organized into subgroups of retinal cells. The majority of horizontal and amacrine cell genes are higher expressed in OVB-Organoids compared to early brain organoids. Likewise, genes typical of bipolar cells are highly up-regulated in OVB-Organoids, whereas genes more generally and not exclusively expressed in Müller Glia cells, progenitor cells and retinal ganglion cells are slightly up-regulated in OVB-Organoids compared to early brain organoids.

**Figure S7.**
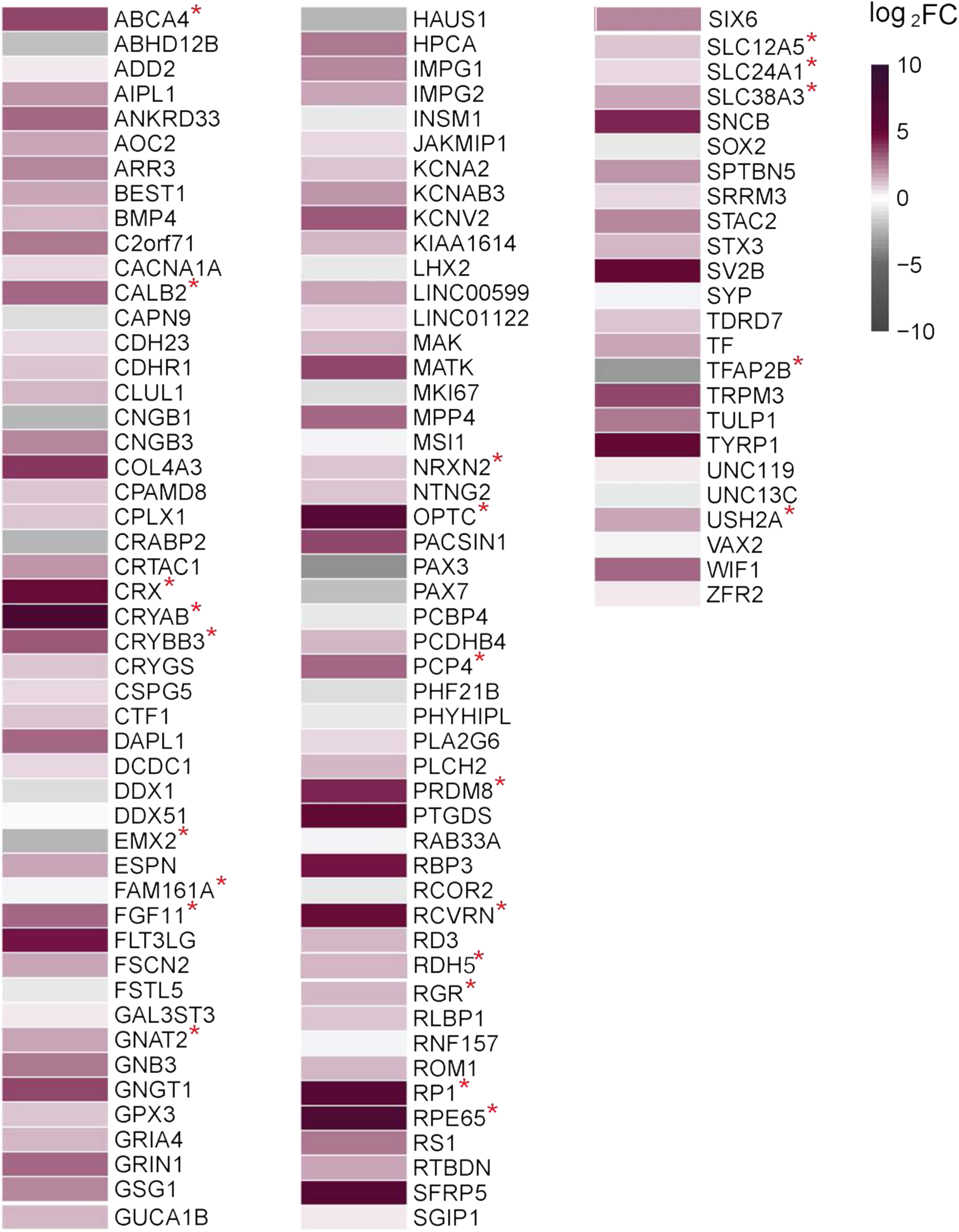
Expression profiling of OVB-Organoids shows compound eye development and optic nerve formation signatures. RNA-Seq results of early brain organoids with primitive eye fields compared to OVB-Organoids. Summary of highly expressed typical eye genes, such as CRX, PCP4, RCVRN (Recoverin), RPE65 (RPE cell marker) TFAP2B (amacrine marker). Red asterisks mark genes expressed in lens (CRYAB, CRYBB3) and Opticin (OPTC), a gene highly expressed in the vitreous humor of the eye, the cornea, ciliary body, optic nerve, choroid, iris, and retina. Besides, genes involved in retinal recycling process such as ABCA4, RDH5, RGR, RP1 and RPE65 were also identified. Other genes or compound eye development, optic nerve and synaptogenesis are CALB2 (Calbindin), NRXN2, SLC12A1, A3 and A5 and genes implicated in retinitis pigmentosa such as FAM161 and USH2A.

**Figure S8.**
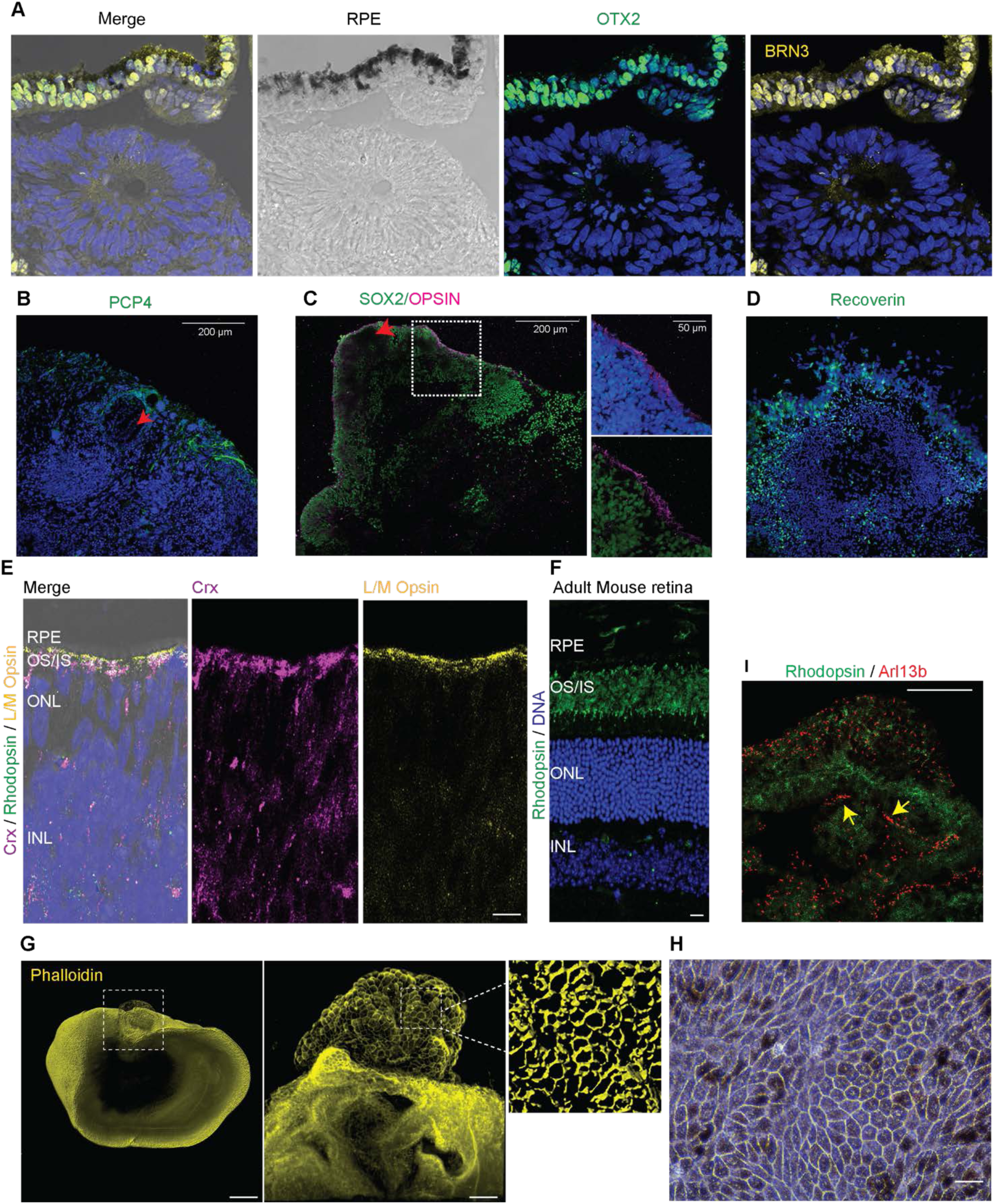
Related to main figure 4. **A.** Transverse section through an optic vesicle of an organoid shows RPE cell layer imaged by transmission light (Dark pigmented). OTX2-positive nuclei (green) co-localized with BRN3 (yellow). Scale bar, 100 µm. N=10 organoids from at least 4 independent batches. Representative image, cell line GM25256. **B-D.** Section through an optic vesicle shows PCP4-positive amacrine cells **(B).** Opsin and recoverin antibodies, which specify photoreceptor cells, mark the infrequent presence developing photoreceptor cells **(C-D)**. Red arrow marks the location of pigmented region within an optic vesicle. Scale bar, 200 µm and 50 µm which are displayed in the respective figures. N=10 organoids from at least 4 independent batches. Representative image, cell line GM25256. **E-F.** Transverse section through an optic vesicle of an organoid shows immature neural retina beginning from RPE cell layer imaged by transmission light (top). Unlike mouse retina **(F)**, OVB-Organoids did not show distinct layers. Cone-rod homeobox protein (CRX), marks a layer adjacent to RPE. This region is also immunoreactive for L/M-opsin. N=10 organoids from at least 4 independent batches. **G.** Surface rendered image of an OVB-Organoid stained with Phalloidin (Yellow). The square at the apex (left panel) highlights the optic vesicle, which is magnified in the middle panel. The right panel is a 3D construction of the RPE within the optic vesicle. Scale bars, 200 µm (left panel), 50 µm (middle panel), 5 µm (right panel). Phalloidin failed to penetrate the inner core of the organoid (left panel). Representative image, cell line GM25256**. C.** Adherent RPE cells dissociated from an optic vesicle form a monolayer RPE sheet displaying typical honeycomb shaped cells filled with pigments. Scale bar, 10 µm. Representative image, cell line IMR-90. **H.** RPE cells plated after disassociation of the pigmented region from an OVB-Organoid. Scale bar, 10 µm. **I.** Immunostaining through an optic vesicle shows Arl13B-positive (red) primary cilia in the pigmented RPE cells (Upper region). Note that pigmented region strongly hinders the visualization of diffused Rhodopsin immunoreactivity. Serially organized primary cilia in a region of Rhodopsin-positive cells (green) below the pigmented cells (yellow arrows). At right, another example with a magnified view. Scale bar, 50 µm. Right image shows a representative image of a second organoid. Scale bar 10 µm. N=11 organoids from 3 independent batches. Representative image, cell line IMR-90.

**Figure S9.**
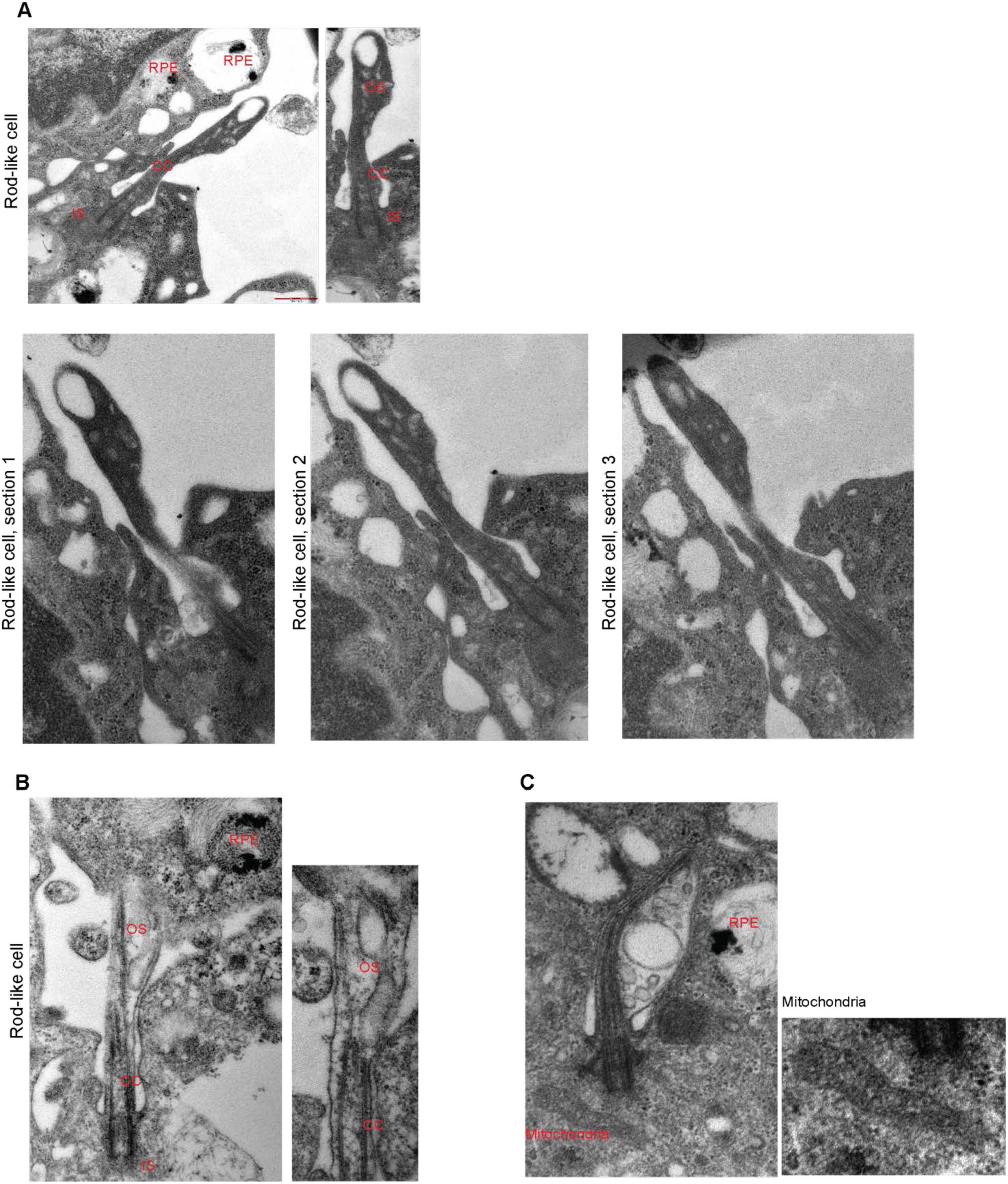
Related to main figure 4. **A.** Serial sectioning and transmission EM image of a developing photoreceptor rod cell. Left panel: The outer segment (OS) is bulging out as an extension at the tip of the connecting cilium (CC). The inner segment (IS) is partly shown and contains the cell body. The photoreceptor cell is neighbored by a retinal pigment epithelial (RPE) cell containing dark pigment granules. Right panel: The outer segment (OS) is filled with vesicles that will later fuse to OS discs. The outer segment appears to develop through an enlargement of the distal part. These longitudinal serial sections additionally showed the process of vesicle formation restricted to only one side of the cilium that is opposite to the side where the axoneme is located. This type of asymmetrical development is a key feature of outer segment differentiation in rod cells of rats(De Robertis, 1956; Kuwabara and Weidman, 1974; Weidman and Kuwabara, 1969) and cat(Tokuyasu and Yamada, 1959). Scale bar, 500 nm. Representative images, cell line IMR-90. **B.** Transmission EM image of another developing photoreceptor rod cell. **C.** Transmission EM image of a cone-like developing photoreceptor cell having a mitochondrion at its base.

**Figure S10.**
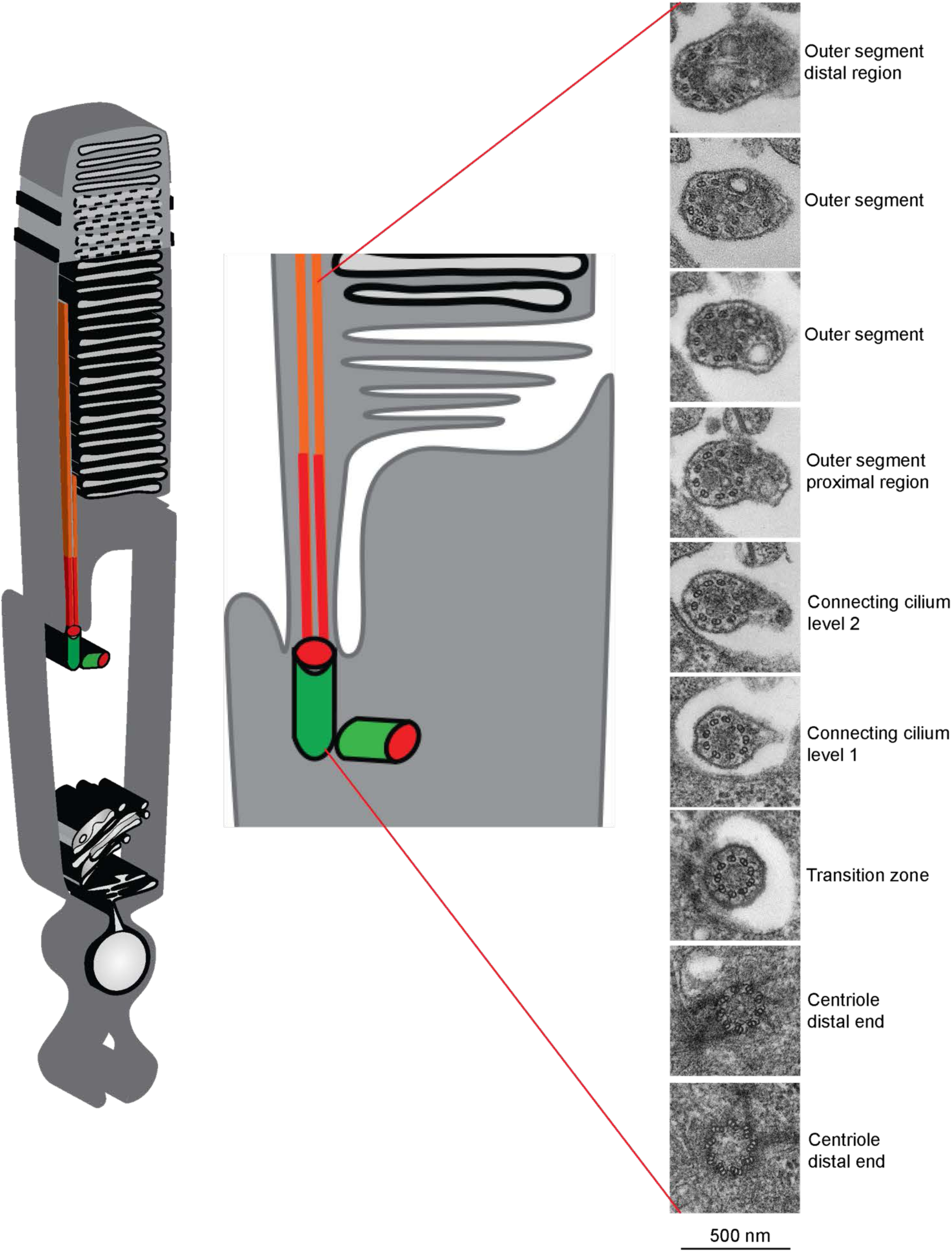
Related to main figure 4. **B.** Serial cross-sections of a developing rod outer segment. The schematic on the left shows a rod cell from the inner segment with nucleus, centrioles (green), connecting cilium (red) and outer segment with discs for photoreception. Middle schematic shows the magnified region from centrioles (green) to outer segment distal region in the longitudinal section for orientation. The right panel shows serial cross-sections through the entire length of the connecting cilium and outer segment. These cross-sections identify that membranous vesicles are indeed asymmetrically distributed to only one side of the cilium. At the bottom, a typical basal body consisting of triplet microtubules with appendages and Y-linkers is seen. At the distal part of the connecting cilium, a small bulge appeared lateral to the axoneme, which continued to grow more significant in the successive sections. More distally, numerous vesicles and flattened cisternae were found within the lateral expansion resembling the pattern of outer segment formation (Obata and Usukura, 1992). Scale bar, 500 nm. Representative images, cell line IMR-90.

**Figure S11.**
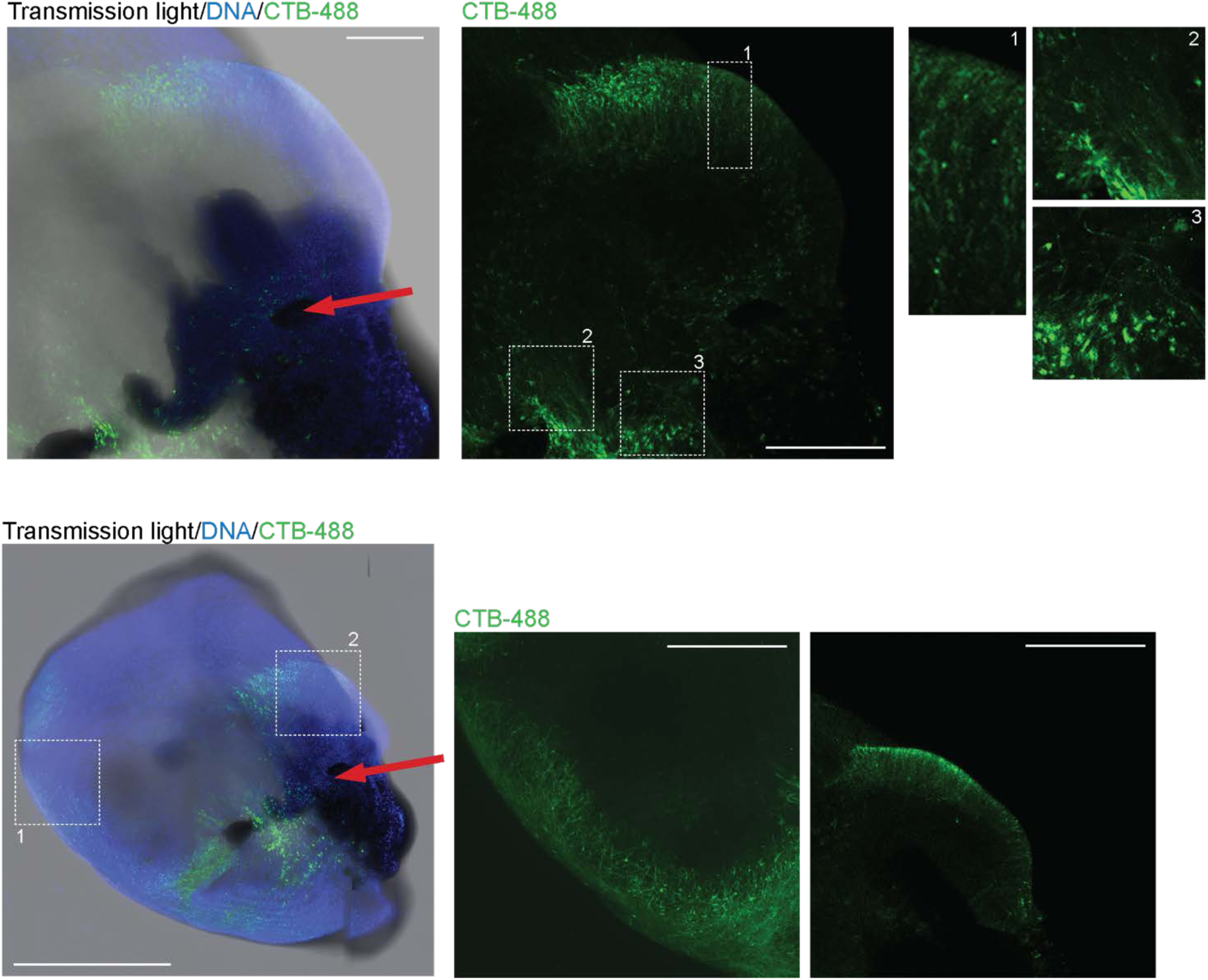
Related to main figure 5. Day 60 old OVB-Organoid was injected with CTB-488 directly into an optic vesicle, and exposed to light for 10 min, cultured for 24 hours and then PFA fixed. Following tissue-clearing, the specimens were imaged. The image on left shows merged transmission light with optic vesicle and dark pigmented area at the injection site (red arrow), nuclear staining (blue) and CTB-488 (green). The middle panel shows the tracing of neuronal pathways by using live dye CTB-488. Right panel 1 shows magnified retinal area, the region of photosensitive neurons that firstly were active upon light stimulation and then transmitted electrical signals and the CTB-488 to next neuronal layers. Right panel 2 and 3 show magnified areas of deeper neuronal layers receiving CTB-488 and thus an electric signal from the photosensitive retinal neurons. Scale bars, 500 µm (left panel), 200 µm (middle and right panel). Bottom panel shows control organoids, which were not exposed to light. Note that there is no apparent CTB uptake or projections are seen. Representative images are given. N=12 from 4 independent batches. Representative images, cell line F14536.2.

**Figure S12.**
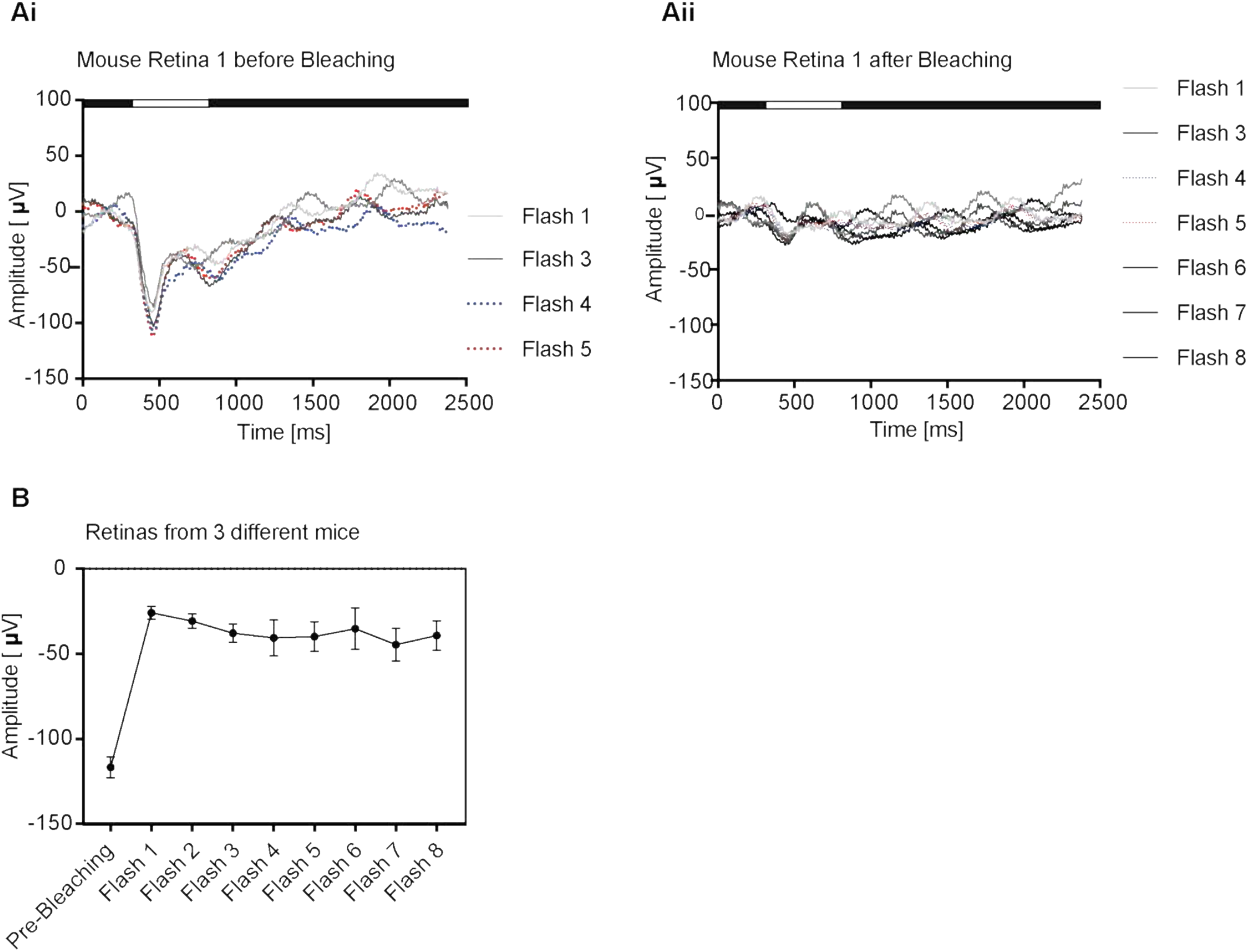
Related to main figure 7. Photic stress experiment with isolated mouse retina. **A.** Mouse retina 1 light response to 200.000 mlux light flashes of 500 ms duration. The recording was done for 5 flashes with each 3 min break (**i.)** The same mouse retina 1 after photic stress by bright light (4600 lux for 10 min with the infrared filter between retina and bright light source) is showing no light response to 200.000 mlux light flashes even after 24 min of recovery **(ii).** **B.** Statistical summary of photic stress experiment with retinas from three different mice. N = 3 independent experiments; One-way ANOVA data followed by Tukeýs multiple comparisons test. Error bars, +/-SEM.

## Supplementary movies

**Movie 1: Imaging of RPE cells within the optic cups after tissue clearing**

Representative movie of F-actin stained (Phalloidin, yellow) RPE shows distinct honeycomb-like architecture. The surface rendered 3D reconstruction model of the honeycomb structure is shown at 00:14 min for improved structural details. Scale bar, 200µm. The representative movie, cell line GM25256.

**Movie 2: Microinjection of Cholera toxin B (CTB-488 and CTB-647) to label axon-like projections**

Representative movie of volume-rendered optic cups microinjected with CTB-488 and CTB-647 at two distinct sites. The injection sites are marked with arrows (00:03 min). Individual channels (GFP and Cy5) have been shown to highlight the distribution of injected toxins. The merging of optic tracts is shown by the strong co-localization of axons that are labeled by both CTB-488 and CTB-647. Scale bar, 200µm. The representative movie, cell line IMR-90.

## Methods & Material

### Human iPS cell maintenance

Human iPS cell lines IMR90, GM25256, F13535.1, F14536.2 and Crx-iPS were cultured in mTeSR1 medium (Stem cell technologies) on Matrigel (Corning) coated cell culture dishes at 37°C and 5% CO_2_ and routinely checked for mycoplasma contamination with the MycoAlert Kit (Lonza). Cells were dissociated to small aggregates with ReLeSR (Stem cell technologies) every 5-7 days and splitted 1:5 onto fresh Matrigel coated dishes. Unless otherwise stated, most analyses shown were done with organoids derived from IMR90 and GM25256.

### Neural induction

It is important to start differentiation with a well-maintained human iPS cell culture. Cells were splitted once after thawing, mycoplasma tested and checked for appropriate stem cell morphology. Before inducing neural differentiation, iPS cells were dissociated to single cells. Therefore, iPS cells were once carefully washed with pre-warmed DMEM/F12 (Gibco) and then treated for 5 min. at 37°C with Accutase (Sigma-Aldrich). After cell collection, centrifuging and counting, cells were diluted in neural induction medium (NIM, Stem cell technologies) plus Rock-inhibitor (10 µM, Y-27632 2HCl, Selleck Chem) at a concentration of 1×10^5^ cells per ml. 100 µl of this cell suspension was given into each well of a 96-well v-bottom low adherent plate and incubated at 37°C and 5% CO_2_. Half of the medium was changed every day for 5 days until neurospheres were formed.

### Neurospheres in petri dishes and spinner flasks

At day 5 of neural induction, the neurospheres were collected carefully with a cut 100 µl pipette tip and transferred to a 100 mm petri dish containing “neurosphere medium” (Method table 1). Additionally, 0.1% Matrigel was added to the medium and neurospheres were incubated for 5 days at 37°C and 5% CO_2_. At day 3, 2 ml of “OVB-Organoid medium” (Method table 2) containing retinol acetate was added. At day 5, neurospheres were transferred to spinner flasks containing 100 ml “OVB-Organoid medium” (Table 2) to let the neurospheres developing to OVB-Organoid. Retinol acetate was added at different concentrations ranging from 0 to 120 nM, but best results (highest yield of OVB-Organoid) were obtained with a concentration of 60 nM retinol acetate. OVB-Organoid were collected for experiments at day 20, 30 and 50. For each cell line, we performed at least 3-4 rounds (batches) of OVB-Organoid generation. Each batch comprises of starting from thawing iPSCs until generating OVB-Organoid. Morphological and functional OVB-Organoid were easily identified by visibly pigmented areas and used for further analyses.

### Immunofluorescence microscopy

OVB-Organoid were collected at day 20, 30 or day 50 and fixed in 4% PFA (Merck Millipore) for 30 min at room temperature (RT) and transferred into new 24-well plates containing PBS-Glycine (0.225 g Glycine per 100 ml PBS). PBS-Glycine was removed, and 30% Sucrose (Merck Millipore) was added to dehydrate the tissues over night at 4°C. Next day, organoids were embedded into Tissue-Tek® Cryomold® Cryomolds (Science services) containing Tissue-Tek® O.C.T.™ Compound solution (Science Services) and immediately stored at −80°C for minimum 24 hours. Frozen specimens were embedded and cryo-sectioned at 12 µm thickness with Cryostat (Leica CM3050 S) and placed onto Poly-L-lysine (Sigma-Aldrich) pre-coated Superfrost glass slides (0.01% PLL diluted in distilled H_2_O and coated for 5 min at RT, subsequently dried 3 hours at RT) (Thermo Scientific). Slides with organoid sections were dried for 1 hour at RT and then kept for long-term storage at −80°C.

Prior to immunofluorescent staining procedures, we thawed the slides with organoid sections for 30 min at room temperature, washed 2 x 3 min with PBS-Glycine (0.225 g per 100 ml) and permeabilized for 15 min in 0.5% Triton X100/0.1% Tween (Sigma Aldrich) in PBS solution at RT, and washed again 2x 3 min with PBS-Glycine. Subsequently sections were blocked with blocking solution (0.5% Fish gelatin in PBS) for 1 hour at RT. Primary antibodies were diluted in blocking solution and sections were incubated with primary antibodies for 2 hours at RT. After 3x washing with blocking solution, corresponding secondary antibodies (goat-anti mouse Alexa 488 or 647; donkey-anti rabbit Alexa 488, 594 or 647 donkey-anti sheep 647 or donkey-anti goat Alexa 594) were diluted in blocking solution (1:500) and incubated for 1 hour at RT protected from light (Method table 4 with antibody details). Finally, sections were stained with nuclear dye Höchst 33342 (Thermofisher), washed 2x 3 min with blocking solution, and after a final wash with distilled water briefly air-dried and then mounted with Mowiol (Sigma Aldrich) and glass coverslips. Slides were dried for one hour at RT and then stored at 4°C in dark until imaging.

### scRNA-sequencing and data analysis

For scRNA library construction, we made use of the Chromium™ Single Cell 3’ Reagent Kits v3. Single nucleus suspensions in 1XPBS containing 0,04% BSA (700-1200 nuclei/µL concentration) have been checked for viability, debris and cell aggregates. To achieve single cell resolution, the cells are delivered at a limiting dilution, such that the majority (∼90-99%) of generated GEMs contains no cell, while the remainder largely contain a single cell. Because of the complex composition of organoids, we aim to target 2.000-3.000 cells per sample. Upon dissolution of the Single Cell 3’ Gel Bead in a GEM, primers containing (i) an Illumina R1 sequence (read 1 sequencing primer), (ii) a 16 bp 10x Barcode, (iii) a 12 bp Unique Molecular Identifier (UMI) and (iv) a poly-dT primer sequence are released and mixed with cell lysate and Master Mix. Incubation of the GEMs then produces barcoded, full-length cDNA from poly-adenylated mRNA. After incubation, the GEMs are broken, and the pooled fractions are recovered. Silane magnetic beads are used to remove leftover biochemical reagents and primers from the post GEM reaction mixture. Full-length, barcoded cDNA is then amplified by PCR to generate sufficient mass for library construction. Enzymatic Fragmentation and Size Selection are used to optimize the cDNA amplicon size prior to library construction. R1 (read 1 primer sequence) are added to the molecules during GEM incubation. P5, P7, a sample index and R2 (read 2 primer sequence) are added during library construction via End Repair, A-tailing, Adaptor Ligation and PCR. The final libraries contain the P5 and P7 primers used in Illumina bridge amplification. A Single Cell 3’ Library comprises standard Illumina paired-end constructs which begin and end with P5 and P7. We allocated Illumina NovaSeq6000 flowcells to sequence with the first read 28nt (cell specific barcode and UMI) and generate with the second read 90nt 3’mRNA transcriptome data. Using the v3 version of chemistry, 25kreads/nucleus are sufficient to allow comprehensive scRNA analysis.Single cell RNA-seq of five batches of organoids was performed using the 10× Chromium pipeline.

Reads were mapped to the human genome (GRCh38), and count matrices were generated using CellRanger v3.1.0. Python notebooks were used to analyze the count matrices, by custom functions and the package Scanpy v1.6.0 (Wolf et al., 2018). Two matrices from organoids at day 30 and three from organoids at day 60 were concatenated. Concatenated matrices were processed with filtering steps to obtain high-quality cells. Low-quality cells were filtered out if they had fewer than 1000 or more than 3000 expressed genes, or more than 0.1% of reads mapping to mitochondrial genes. Expression values were then normalized to 10^4^ and log transformed. The top 1000 highly variable genes were then selected for mutual nearest network (mnn) correction of the counts and dimensionality reduction (Haghverdi et al., 2018). A batch balanced nearest network (bbknn) was then computed on the first 50 principal components (Polański et al., 2019). Such network has been used for visualization of reduced dimensions of the data by UMAP and Leiden clustering (resolution = 0.8). Marker genes characterizing each cluster were calculated by Student’s *t*-test between each group and all the other cells. Annotation of clusters was performed by inspection of marker genes and plotting of known cell type specific genes. Pseudotime in d30 dataset was computed by the diffusion pseudotime (DPT) algorithm setting the stem cell-like cluster as root (Haghverdi et al., 2016).

### Microinjection and tissue clearing

For optic tract experiments, we injected CTB-488 or CTB-647 (Invitrogen) (1:1 (v/v) mixed with DMEM/F12 comprising a total volume of 3 µl) into each of the OVB-Organoid recognized by the pigmented areas within the surrounding whitish tissue. We used a micromanipulator (Märzäuser, Wetzlar) for injection. Injected organoids were placed into wells of a 24-well plate in OVB-Organoid medium and then exposed to normal room light for 10 min. After exposure to light, the organoids were further cultured at 37°C and 5% CO2 for 24 hours before fixation with 4% paraformaldehyde (PFA) for 30 min at RT.

Tissue clearing was performed as previously described(Masselink et al., 2019). In brief, PFA fixed organoids were dehydrated sequentially in graded ethanol (EtOH) series of 30% EtOH (vol/vol), 50% EtOH (vol/vol), 75% EtOH (vol/vol) and 100% EtOH (vol/vol) (each step for 30 min at 4***°***C in a gently shaking tube 1ml tube) and subsequent tissue clearing was performed with Ethyl cinnamate (ECI; Sigma Aldrich) for approximately 30 min at room temperature. Tissue cleared organoids were then placed into cover slip bottom µ-slides (Ibidi) and stored at 4°C until 3D imaging.

### 3D imaging and processing

Images were acquired with confocal microscope (Leica SP8) and converted into .tiff files with Fiji software (ImageJ 1.52i). Images were then processed with Photoshop (Adobe CS5) and copied as .psd files into Adobe Illustrator. Tissue cleared organoids were placed into µ-Slide angiogenesis (Ibidi, Germany) and imaged using Zeiss LSM 880 confocal microscope with EC Plan-Neoflaur 10x/0.3 and Plan-Apochromat 20x/0.8 M27 objectives. The raw files were exported to Imaris x64 version 7.7.1 to perform surface and volume rendering. The files were further processed using Image J, Adobe photoshop CC 2017 and Adobe illustrator CC 2017.

### Electrophysiology: ERG with organoids

OVB-Organoid of day 50 to 60 were transferred to wells of a 12-wellplate each with 2 ml of OVB-Organoid medium and kept in absolute darkness at 37°C and 5% CO2 for 24 hours before electroretinogram (ERG) experiments. For ERG, each organoid was placed under red light in an electrode chamber. The electrode chamber was then placed into the dark chamber of the ERG apparatus and connected to the superfusion system containing optic organoid medium.

For dose dependent light sensitivity, organoids were exposed to 500 milliseconds long flashes of 2000, 20.000 and 200.000 mlux. Each light intensity was repeated three times in a time interval of 3 min. before the next higher light intensity was applied and responses were recorded. Speed of superfusion was 0.5 ml/min. In controls, we used tissues without optic vesicles, which seldom displayed a basal level of response.

For photic stress experiment, OVB-Organoid were superfused with 0.5 ml/min OVB-Organoid and exposed to 500 ms long flashes of 200.000 mlux every 3 min. for adjustment to the system for 15 min. (pre-bright light). Then organoids were desensitized in electrode chamber by bright light exposure with light intensity of 4600 Lux for 10 min. As bright light source served a 150 W white light bulb. Between bulb and electrode chamber was an infrared filter placed to subtract the thermal impact of the bright light source. After 10 min. of bright light exposure, organoids were allowed to regenerate in dark chamber for 30 s before light flashing (500 ms, 200.000 mlux) and recording under above mentioned conditions continued. These experiments never included 11-cis-retinal as such at any time point. As negative control, when organoids were regenerated from photic stress usually after 30 min. of flashing and recording, same organoids were treated under same conditions again except bright light exposure was replaced by 0 lux.

For a third experiment, organoids were superfused in optic organoid medium with higher speed of 2 ml/min. After 36 min of equilibration time, organoids got adjusted to the higher superfusion speed and responded with a stable electrical signal of −200 mV as response to flashes of 200.000 mlux and 500 ms. For isolating the A-wave by blocking signal transduction from photoreceptors to next the retinal cell layers, 10 mM aspartate was added to the medium after the equilibration time and electric response was continuously recorded. Finally, aspartate was washed out by adding optic organoid medium without aspartate into the system while recording of electric signals as response to flashes.

### Electrophysiology: ERG with isolated mouse retina

Mice were dark-adapted overnight, sacrificed by cervical dislocation under a dim red light and the eyes were extirpated immediately. Enucleated eyes were protected from light and transferred into carbogen-saturated (95 % O_2_ / 5 % CO_2_) Ames medium and into the recording chamber as described previously(Albanna et al., 2017). The b-wave amplitude was measured from the trough of the a-wave to the peak of the b-wave. For each experiment, a new murine retina was transferred to the recording chamber. The institutional and governmental committees on animal care approved the animal experimentation described in the text (Landesamt für Natur, Umwelt und Verbraucherschutz (LANUV) Nordrhein–Westfalen, Recklinghausen, Germany; 84–02.04.2016.A4555) which was conducted following accepted standards of humane animal care.

### Electrophysiology: Whole cell patch clamp

Living organoids were embedded in 2% low-melt agarose (Invitrogen) and attached to a cutting plate beside a 4% agarose block. Vibratome sectioning was done at room temperature. For patch clamp experiment slices were cut into 350-400 µm sections.

Cells with spontaneous activity were identified before patch clamping by staining with fluo8AM (Abcam) (5µg/ml) for 15 min at room temperature. Overview of spontaneous activity was recorded with a LSM setup (Zeiss 780). Each cell was recorded for 5 to 10 min. in “on-cell”/” cell” attached configuration while the fluorescent signal was recorded as well. Then the cell membrane was broken to perform “whole-cell” configuration for channel currents recording. During whole cell recording, the cell is clamped at a resting potential of −70 mV. Depolarization steps were applied in a 10-mV interval for the current-voltage curves. The recording temperature was approx. 37°C in ACSF (artificial spinal fluid).

### Transmission electron microscopy (TEM)

Organoids of day 50 were washed once in PBS and then fixed in 2.5% Glutaraldehyde and 1.8% Sucrose in PBS over night at 4°C. Samples were rinsed in PBS, post-fixed with 1% osmium tetroxide for 1 hr at 4°C. After fixation, the samples were further dehydrated in a graded ethanol series and embedded in Epon-Araldite resin for 48 hrs at 60°C. Reichert Ultracut E ultramicrotome was used to obtain the Ultrathin sections (40-60nm), further stained with uranyl acetate and lead citrate, and observed with a FEI Tecnai G2 Spirit 255 transmission electron microscope operating at 100 kV.

### Scanning electron microscopy (SEM)

Organoids of day 50 were washed once in PBS and then fixed in 2.5% Glutaraldehyde and 1.8% Sucrose in PBS over night at 4°C. Samples were then rinsed with PBS and post-fixed with 1% osmium tetroxide in PBS for I hr at 4°C. After alcohol dehydration, samples were critical point dried in a Balzers CDP 010. The material was then mounted on aluminum stubs, coated with gold using a Balzers Med 010 and examined with a Philips XL20 scanning electron microscope (SEM) operating at 15 KV.

### RNA isolation, RNA-sequencing and analysis

Organoids (from IMR-90) were washed 1x in cold PBS and then immediately transferred to a fresh RNAse free vessel containing 300 µl Tri-Reagent (Sigma Aldrich, USA) by free-thawing, and total RNA was isolated and DNase-treated using the DirectZol RNA kit (Zymo Research). Approximately 2 μg of total RNA was used to subselect poly(A)+ transcripts and generate strand-specific cDNA libraries (TrueSeq kit; Illumina). These libraries were then sequenced to ∼50 million read pairs each (75-bp reads) on a HiSeq4000 platform (Illumina).

After assessing the sequencing data quality with FastQC, paired-end sequencing reads were mapped with STAR aligner(Dobin et al., 2013) to Hg19 human reference assembly and quantified with featureCounts(Liao et al., 2014). Raw read counts were normalized with RUVSeq(Risso et al., 2014) package in R in order to remove unwanted variation based on replicate samples (RUVs). Data was subsequently analyzed for changes in differential gene expression (DGE) with DESeq2(Love et al., 2014) package. List of 3923 significantly DGE genes (log2FC ±1.5 and padj<=0.05 cutoff) was subjected to affinity propagation clustering based on similarity matrix, with apcluster(Bodenhofer et al., 2011) package. From resulting dendrogram containing 75 clusters, cutoff of k=9 resulted in 9 super clusters which were used for GO term search analysis with clusterProfiler(Yu et al., 2012) Significant DGE genes classified in super-cluster 9 were subjected to gene set enrichment analysis with GSEA (ref 8**; v3.0) using C5 collection of GO gene sets from Molecular Signatures Database(Liberzon et al., 2011)

### RPE isolation and culturing

For isolation of RPEs, OVB-Organoid were grown until day 50-60 and then dissociated with Trypsin (0.05%, 20 min, 37°C) to small cell aggregates. Trypsin was inhibited by 20%-DMEM-FBS and dark aggregates were collected in a separate 15 ml Falcon, and centrifuged at 1000x g. The pellet was resuspended in RPE Medium (Method Table 3) and cells were seeded onto laminin-coated cell culture treated dishes. Cells were cultured approx. for two weeks until large colonies of pigmented cells were grown as monolayer sheets.

### Statistics

Statistical analyses were performed using Graphpad Prim 7 for Mac OS X (Version 7.0e, September 5, 2018).

**Method table 1.**
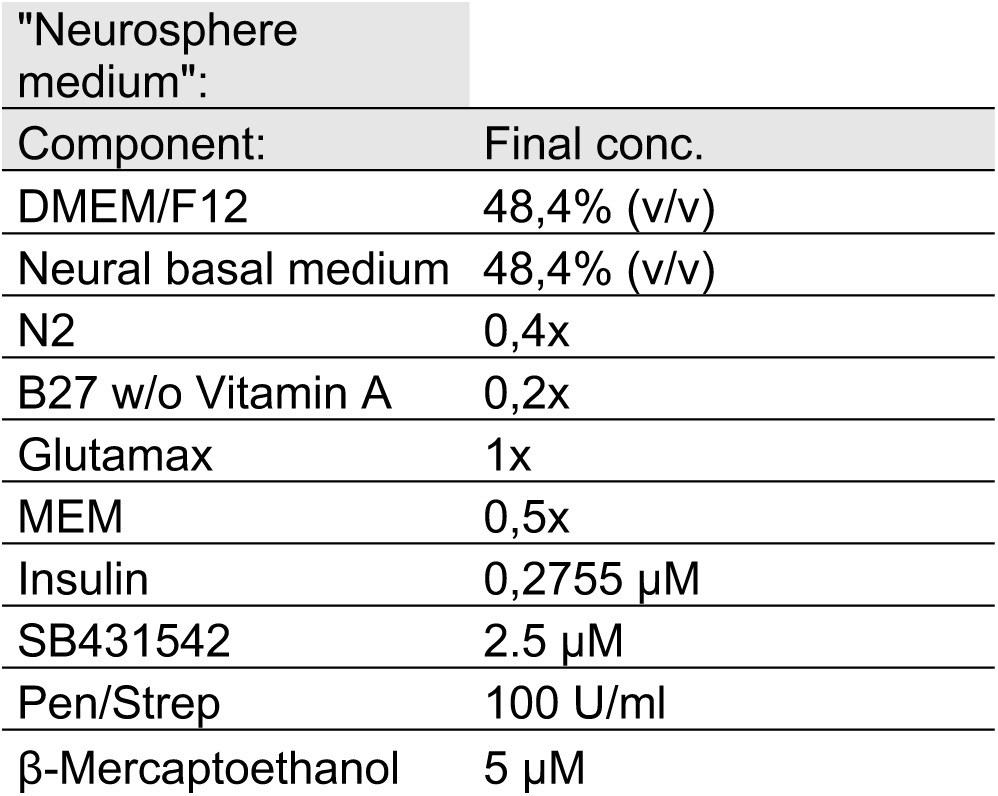

**Method table 2.**

| " OVB-Organoid medium": |  |
| --- | --- |
| Component: | Final conc. |
| DMEM/F12 | 48,4% (v/v) |
| Neural basal medium | 48,4% (v/v) |
| N2 | 0,4x |
| B27 + RA | 0,2x incl. 60 nMol retinol acetate |
| Glutamax | 1x |
| MEM | 0,5x |
| Dorsomorphin | 0.5 µM |
| Insulin | 0,2755 μM |
| SB431542 | 5 μM |
| Pen/Strep | 100 U/ml |
| β-Mercaptoethanol | 5 μM |

**Method table 3.**

| "RPE medium": |  |
| --- | --- |
| Component: | Final conc. |
| DMEM/F12 | 87% (v/v) |
| FBS | 10% (v/v) |
| Glutamax | 1x |
| MEM | 1x |
| Pen/Strep | 100 U/ml and 100 μg/ml |
| β-Mercaptoethanol | 5 μM |

**Method table 4.**
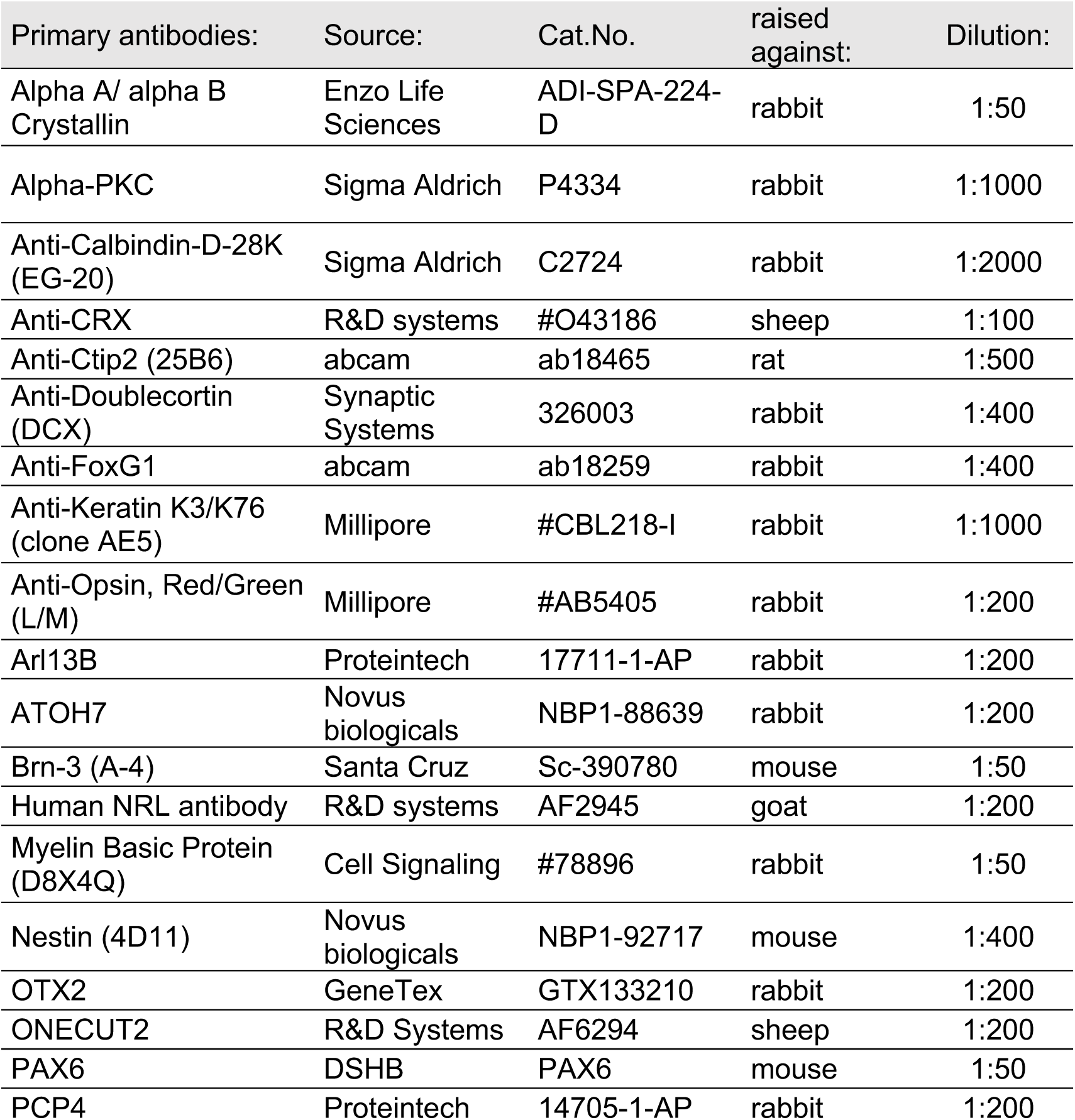
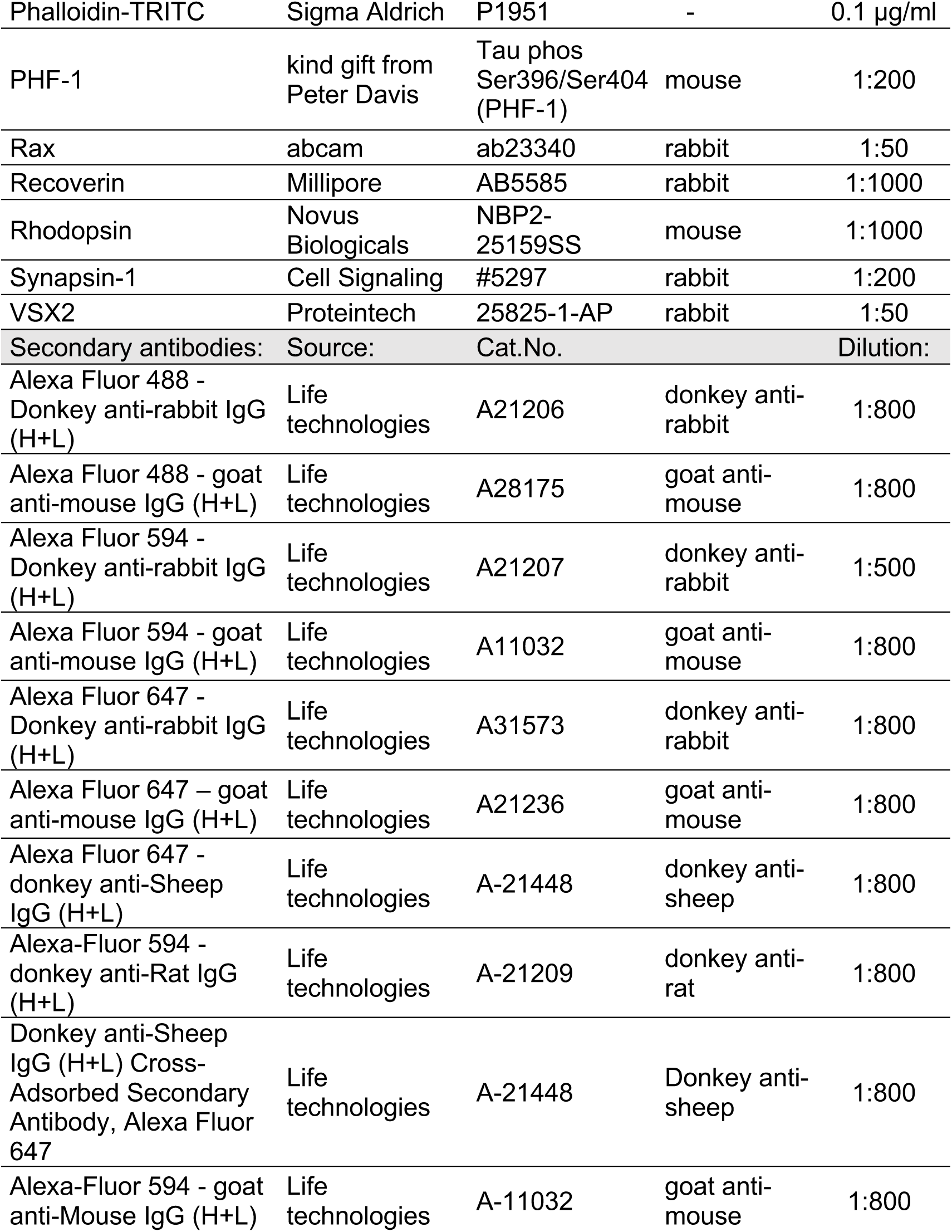

